# INO80 promotes H2A.Z occupancy to regulate cell fate transition in pluripotent stem cells

**DOI:** 10.1101/2020.12.09.418400

**Authors:** Hongyao Yu, Jiajia Wang, Brad Lackford, Brian Bennett, Jian-liang Li, Guang Hu

**Affiliations:** Epigenetics and Stem Cell Biology Laboratory, National Institute of Environmental Health Sciences, Research Triangle Park, North Carolina, 27709, USA; Integrative Bioinformatics Support Group, National Institute of Environmental Health Sciences, Research Triangle Park, North Carolina, 27709, USA

**Author notes:** These authors contributed equally to this work (Hongyao Yu, Jiajia Wang). Correspondence and requests for materials should be addressed to: Hongyao Yu, Ph.D., Guang Hu, Ph.D.

## Abstract

The INO80 chromatin remodeler is involved in many chromatin-dependent cellular functions. However, its role in pluripotency and cell fate transition is not fully defined. We examined the impact of Ino80 deletion in the naïve and primed pluripotent stem cells. We found that Ino80 deletion had minimal effect on self-renewal and gene expression in the naïve state, but led to cellular differentiation and de-repression of developmental genes in the transition toward and maintenance of the primed state. Mechanistically, INO80 pre-marked gene promoters that would adopt the H3K4me3 and H3K27me3 bivalent histone modifications. It promoted H2A.Z occupancy at these future bivalent domains to facilitate H3K27me3 installation and maintenance as well as downstream gene repression. Thus, INO80-dependent H2A.Z occupancy is a critical a licensing step for bivalency and poised gene expression in pluripotent stem cells. Our results uncovered an unexpected function of INO80 in H2A.Z deposition and gene repression, and an epigenetic mechanism by which chromatin remodeling, histone variant and modification coordinately control cell fate.

## Introduction

Pluripotent stem cells can provide important insights to both stem cell and developmental biology. Recent studies showed that there are different pluripotent states that correspond to the sequential developmental stages in embryos(1,2). These states can be captured *in vitro* using defined culture conditions, and they are characterized by distinct developmental potential, gene expression profile, and epigenetic status(3–12). Among them, the naïve pluripotent state is the earliest stage in pluripotency during embryonic development. It is represented by mouse embryonic stem cells (ESCs) cultured in medium containing MEK, GSK3 inhibitors and leukemia inhibitory factor (LIF) (2iL)(3). The primed pluripotent state is a later stage, and is represented by mouse epiblast stem cells (EpiSCs) cultured in medium containing FGF2 and Activin A (FA)(13,14) or FGF2, Activin A and the tankyrase inhibitor XAV939 (FAX)(15,16). These pluripotent stem cell models are valuable tools to investigate the molecular events that regulate pluripotency and early development.

The chromatin of pluripotent stem cells is characterized by unique epigenetic features that contribute to their special developmental potentials. Of particular interest, the pluripotent chromatin contains regions that are decorated by both the active histone H3 lysine 4 trimethylation (H3K4me3) and repressive H3 lysine 27 trimethylation marks (H3K27me3)(17). These bivalent chromatin domains are enriched at the promoters of key developmental genes(17), and they facilitate the formation of specific 3D chromatin conformations(18). The presence of the two apparently contradictory histone marks set downstream developmental genes in a poised state, which contribute to the intricate balance between self-renewal and differentiation(19,20). Bivalency is quickly resolved into either the active or repressive state once the cells differentiate and is therefore can be viewed as a chromatin property that is associated with pluripotency(17). However, how the bivalent domains are formed and regulated remain incompletely understood. The Polycomb Repressive Complex 2 (PRC2) catalyzes H3K27 methylation and is essential for bivalency(21). Intriguingly, H3K27me3 can be accurately established *de novo* at bivalent domains when PRC2 activity is restored in PRC2-deficient ESCs(22). Similarly, the bivalent modifications are initially absent from developmental gene promoters in mouse pre-implantation embryos but are soon established after implantation(23,24). Thus, there likely exist additional factors that facilitate the establishment and maintenance of the bivalent domains.

Because of the distinctive chromatin characteristics, it is not surprising that pluripotent stem cells are highly dependent on chromatin regulators such as the chromatin remodeling enzymes(25). The chromatin remodelers use the energy of ATP hydrolysis to regulate chromatin structure and dynamics. They can change nucleosome compositions, or remove, exchange, and slide nucleosomes, and thereby play critical roles in a wide range of chromatin-dependent functions(26). The INO80 chromatin remodeling complex belongs to the INO80 sub-family of chromatin remodelers(27,28). It has DNA-dependent ATPase activity and catalyzes ATP-dependent nucleosome sliding and spacing(27,29), as well as the exchange of the histone variant H2A.Z(30,31). INO80 has been implicated in many cellular and physiological processes such as stem cell maintenance, embryogenesis, germ cell development, neurological diseases, and cancer(32–36). In mouse ESCs cultured in serum and LIF, INO80 occupies pluripotency gene promoters and facilitates downstream gene activation(33). In mice, INO80 promotes proximal-distal axis establishment and its deletion led to early embryonic lethality(32,35). Therefore, INO80 plays important roles in pluripotency and early development.

As earlier studies relied on ESCs cultured in heterogenous conditions, the role of INO80 in defined pluripotent states has not been carefully investigated. Here, we set out to examine the impact of *Ino80* deletion on the naïve and primed pluripotent state. We found that INO80 is selectively required in the primed but not the naïve state. It promotes H2A.Z occupancy to facilitate the establishment and maintenance of the bivalent domains at developmental genes, in order to keep them in the poised state. Our findings uncovered a critical step in the formation and regulation of the bivalent chromatin structure in pluripotent stem cells. As both INO80 and H2A.Z are heavily involved in development and disease, we propose that the INO80-H2A.Z axis may have similar functions during other cell fate transitions.

## Results

### INO80 is selectively required for the maintenance of the primed pluripotent state

To investigate the role of INO80 in pluripotent stem cells, we generated an inducible deletion mouse ESC line for the core ATPase INO80 (*Ino80*-cKO ESCs, Supplementary Fig. 1a). The Ino80-cKO cells have a normal karyotype (Supplementary Fig. 1b) and show efficient deletion of *Ino80* at both the mRNA and protein levels upon 4-hydroxytamoxifen (4-OHT) treatment (Supplementary Fig. 1c-d). Moreover, chromatin immunoprecipitation followed by high throughput sequencing (ChIP-seq) using an antibody that recognizes the C-terminus of INO80 confirmed the loss of INO80 genomic occupancy in the deletion cells (Supplementary Fig. 1e).

In order to study the naïve and primed pluripotent states, we cultured ESCs in the 2i/LIF (2iL) or FGF2/Activin-A/XAV-939 (FAX) medium as previously described(3,4,8,9,16). We were also able to induce the transition from the naïve to the primed state by switching the cells from the 2iL to FGF2/Activin-A (FA) medium (Fig. 1a)(37). With these *in vitro* culture models, we set out to systematically dissect the molecular function of INO80 in different pluripotent states.

**Figure 1.**
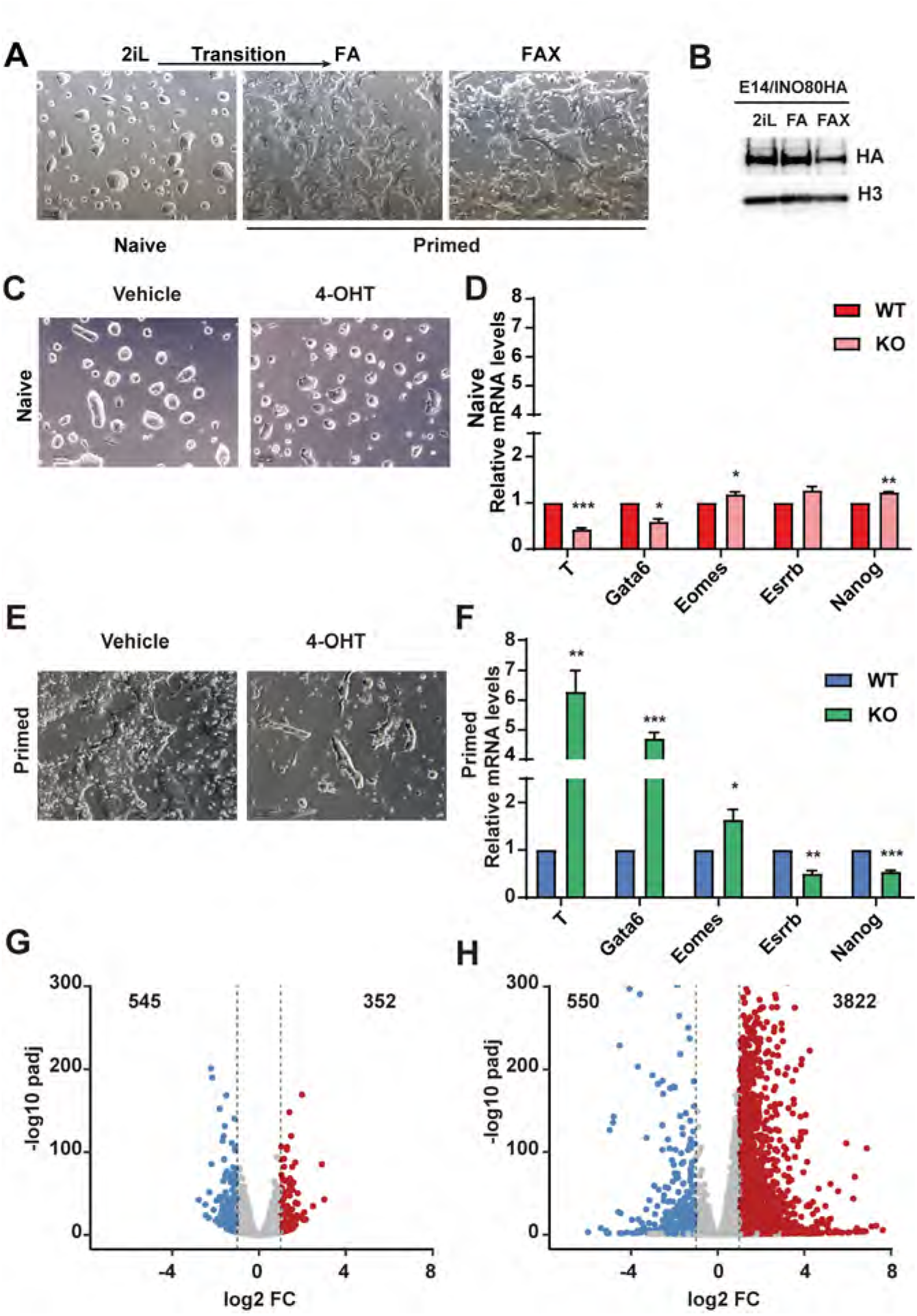
INO80 is selectively required in the primed pluripotent state. **(a)** Culture model for the naïve and primed pluripotent state. Mouse ESCs were cultured in 2iL for the naïve state, switched to FA to induce the transition toward the primed state, and maintained in FAX for the primed state. Scale bar = 100 μm. **(b)** INO80 expression in different pluripotent states. *Ino80-HA* knock-in ESCs were cultured in 2iL, FA, or FAX, and INO80 expression was detected by western blot using the HA antibody. Histone H3 was blotted as the loading control. **(c-d)** *Ino80* deletion in the naïve state. *Ino80*-cKO ESCs were cultured in 2iL and treated with DMSO or 4-OHT for 2 days to induce *Ino80* deletion. Cell morphology (c) and lineage marker expression (d) were determined by imaging and RT-qPCR. Scale bar = 100 μm. Relative mRNA level was normalized with *Gapdh* and plotted as mean ± SEM. p-values were calculated by student *t*-test: * <0.05, ** <0.01, *** <0.001. **(e-f)** *Ino80* deletion in the primed state. *Ino80*-cKO ESCs were cultured in FAX and treated with DMSO or 4-OHT for 2 days. Cell morphology (e) and lineage marker expression (f) were determined by imaging and RT-qPCR. **(g-h)** Gene expression changes after *Ino80* deletion in the naïve and primed state. Differentially expressed genes (DEGs) were determined by RNA-seq and highlighted in the volcano plots. Numbers of DEGs were also listed. Blue: down-regulated genes; Red: up-regulated genes.

The INO80 protein was expressed at comparable levels in the naïve state and during the naïve-to-primed transition, but became slightly reduced in the primed state (Fig. 1b). Surprisingly, *Ino80* deletion had no effect on cell morphology (Fig. 1c) and minimal impact on the expression of the pluripotency and differentiation markers in the naïve state (Fig 1d). Furthermore, *Ino80*-null ESCs can be stably maintained for more than 5 passages in 2iL. In the primed state, however, *Ino80* deletion led to differentiation and cell death (Fig. 1e). In addition, the deletion resulted in reduced expression of pluripotency markers such as *Nanog* and *Esrrb*, and increased expression of lineage markers such as *Eomes Gata6*, and *T* (Fig. 1f). Together, these results indicated that Ino80 is dispensable for the naïve state but required for the primed pluripotent state.

To understand why INO80 is selectively required in the primed state, we carried out RNA-seq to examine the impact of *Ino80* deletion on gene expression. We found that *Ino80* deletion in the naïve state only led to modest changes in the transcriptome (Fig. 1g), consistent with the cellular phenotype. However, its deletion in the primed state resulted in much more prominent perturbations, as shown by both the number and magnitude of differentially expressed genes (Fig. 1h). It also led to a strong bias in gene expression changes, with 3822 up-regulated but only 550 down-regulated genes (Fig. 1h). This is consistent with the phenotypic observation (Fig. 1e) and further suggested that INO80 primarily represses gene expression in the primed state.

### INO80 occupies bivalent gene promoters

To investigate how INO80 regulates gene expression, we examined its genomic occupancy by ChIP-seq. We made the following improvements to the INO80 ChIP-seq strategy. First, we generated a knock-in ESC line by inserting the HA-tag in the C-terminus of the endogenous *Ino80* and carried out INO80 ChIP-seq with the HA antibody (Supplementary Fig. 2a). The HA-antibody ChIP produced a larger number of peaks (Supplementary Fig. 2b), and improved signal-to-noise ratio (Supplementary Fig. 2c) over the INO80-antibody ChIP used in previous works(33,34). Second, we used the MNase digestion instead of sonication for chromatin fragmentation. In addition to the peaks identified by sonication, MNase ChIP-seq captured many more INO80-bound peaks (Supplementary Fig. 2d-e). It is worth noting that these additional peaks were located in regions with lower H3K4me3 but higher H3k27me3 signals (Supplementary Fig. 2f), suggesting that they may be less accessible and less efficiently recovered in sonication-based ChIP-seq.

With the optimized ChIP-seq procedure, we determined the genomic occupancy of INO80 in both the naïve and primed pluripotent state. We found that INO80 occupies a large number of genomic regions in both states, and preferentially binds near the transcription start sites (TSSs) in the primed state (Supplementary Fig. 2g). When intercepted with the RNA-seq data from the *Ino80* deletion cells, INO80 genomic occupancy was found near many of the differentially expressed genes (Supplementary Fig. 2h), suggesting that INO80 may directly regulate their expression. In agreement with previous reports(33,34), many INO80-bound TSSs were marked by the active histone mark H3K4me3 (Fig. 2a). To our surprise, however, a significant fraction of INO80-bound TSSs were bivalently marked by both H3K4me3 and H3K27me3, especially in the primed state (Fig. 2a). In fact, INO80 occupied the majority of the bivalent promoters in the primed state (Fig. 2b), suggesting that it may play an important role in their regulation. This is consistent with the recent discovery that human INO80 occupies both active and repressive chromatin domains (38).

**Figure 2.**
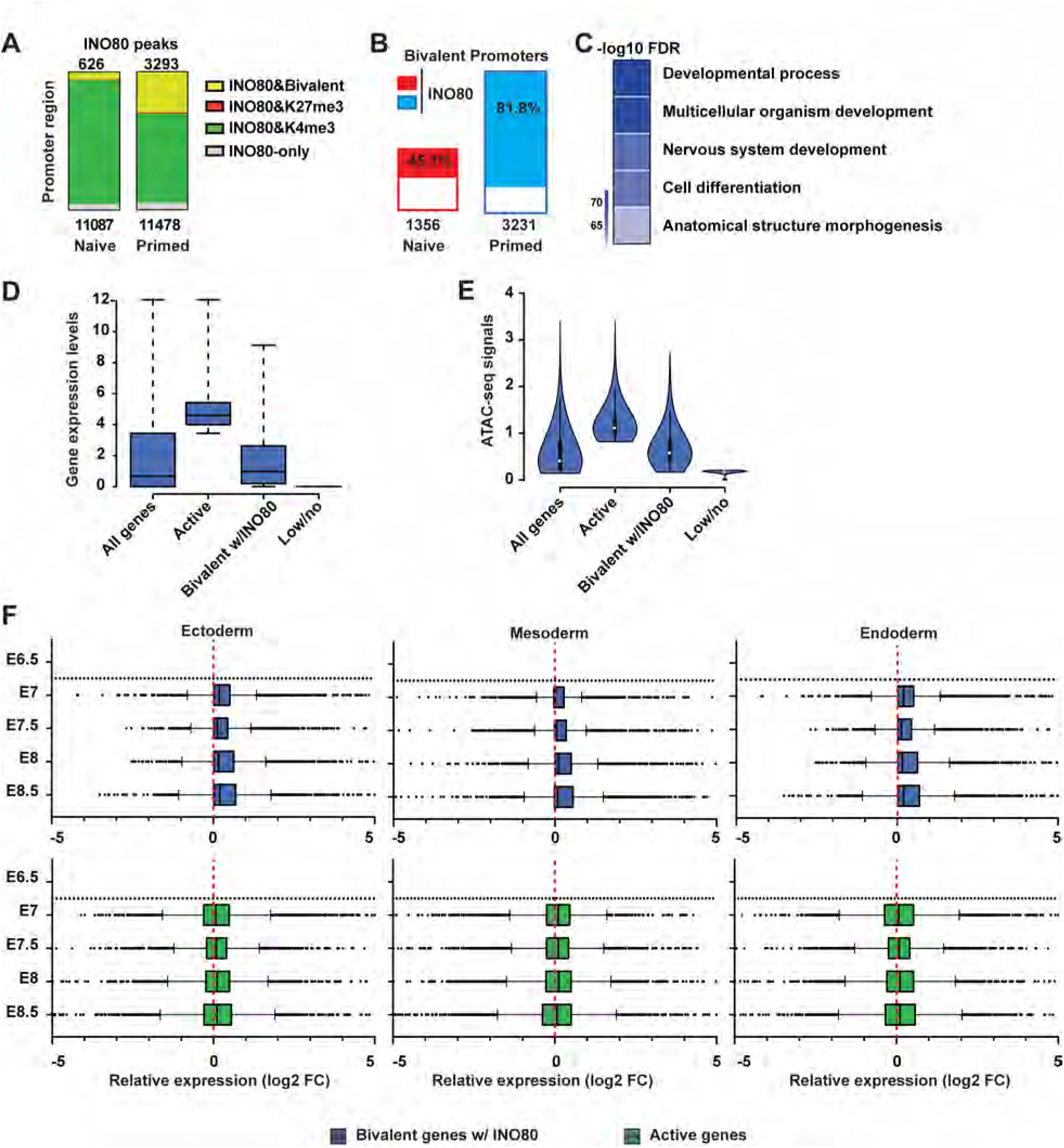
INO80 occupies bivalent gene promoters. **(a)** H3K4me3, H3K27me3 and bivalent domains in INO80 occupied promoters in the naïve and primed state. Number of total INO80 peaks (bottom) and INO80 peaks with bivalent marks (top) in the promoter region were listed. **(b)** Percentage of bivalent promoters occupied by INO80 in the naïve and primed state. Number of total bivalent promoters were listed at the bottom. **(c)** Gene Ontology analysis of INO80 bound bivalent genes in primed state. Selected top categories were shown and the complete list of enriched categories can be found in Supplementary Fig. 2i. **(d)** Box plots to show the relative expression of all genes, active genes (top 25%), INO80-bound bivalent genes, and lowly or not-expressed genes (bottom 25%) in the primed state. Gene expression was determined by RNA-seq and calculated as Log2 (RPKM+ 1). **(e)** Violin plot to show the chromatin accessibility of all genes, active genes (top 25%), INO80-bound bivalent genes, and lowly or not-expressed genes (bottom 25%) in the primed state. Chromatin accessibility was determined by ATAC-seq. **(f)** Box plots to show relative gene expression changes of active and INO80-bound bivalent genes as in D during mouse embryonic development from E6.5-E8.5. Gene expression changes between Ectoderm, Mesoderm, or Endoderm cells and E6.5 epiblast cells were calculated from published single cell RNA-seq data.

### INO80 is required for the establishment and maintenance of the bivalent gene promoters

To test whether and how INO80 regulates bivalent genes, we examined the behavior of the INO80-bound bivalent genes in the presence and absence of *Ino80* in the primed state. We found that the INO80-bound bivalent genes were highly enriched for those that are involved in development (Fig. 2c and supplementary Fig. 2i). They were expressed at low to moderate levels in comparison to highly- and lowly/un-expressed genes (Fig. 2d), and their promoters displayed low to moderate chromatin accessibilities (Fig. 2e). In addition, they were generally up-regulated during mouse embryonic development from the epiblast cells at E6.5 to the three main germ layers at E8.5, while active genes did not show the same trend (Fig. 2f). Thus, INO80-bound bivalent genes appear to be poised in pluripotent cells and activated at the onset of differentiation and lineage commitment. Importantly, the INO80-bound bivalent genes became derepressed upon *Ino80* deletion. A significant fraction of them were up-regulated in the primed state (Fig. 3a-b), and many of them also showed premature or excessive activation during embryoid body differentiation (Fig. 3c). Therefore, Ino80 is required for the poised expression of bivalent genes and prevents them from temporal-spatially mis-regulated activation.

**Figure 3.**
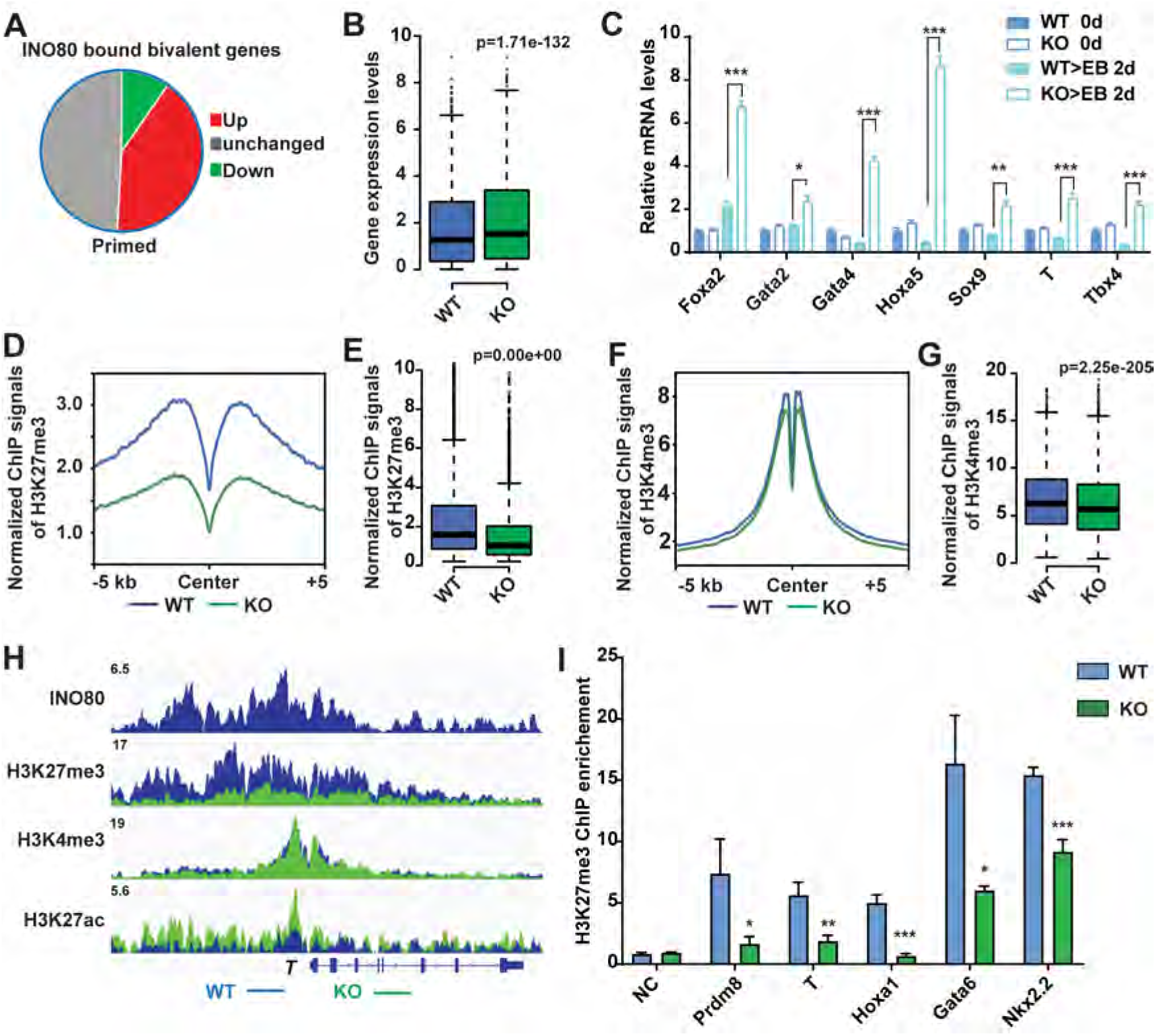
INO80 is required for H3K27me3 occupancy at the bivalent promoters. **(a)** DEGs in INO80-bound bivalent genes in the primed state. **(b)** Box plots to show changes in the expression of INO80-bound bivalent genes. p-value was calculated by the Wilcoxon signed rank test. **(c)** Expression of selected INO80-bound bivalent genes during EB differentiation. ESCs were cultured in FA and treated with DMSO or 4-OHT to induce *Ino80* deletion for two days, and then aggregated to form EBs for 2 days to initiate differentiation. Relative mRNA expression was determined by RT-qPCR, normalized by *Gapdh*, and plotted as mean ± SEM. p-values were calculated by student *t*-test: * <0.05, ** <0.01, *** <0.001. **(d-e)** H3K27me3 occupancy at INO80-bound bivalent TSSs in WT and *Ino80* deletion cells in the primed state. ESCs were cultured in FAX and treated with DMSO or 4-OHT to induce *Ino80* deletion for two days. H3K27me3 occupancy was determined by ChIP-seq, and normalized ChIP-seq signal was used for metagene (d) and box plots (e). p-value was calculated by Wilcoxon signed rank test. **(f-g)** H3K4me3 occupancy at TSS in WT and *Ino80* deletion cells in the primed state. **(h)**, Genome browser view of INO80, H3K27me3, H3K4me3, and H3K27ac occupancy near T in WT and *Ino80* deletion cells in the primed state. **(i)** H3K27me3 occupancy at representative bivalent gene promoters as determined by ChIP-qPCR. Fold-enrichment was plotted as mean ± SEM. p-values were calculated by student *t*-test: * <0.05, ** <0.01, *** <0.001.

To understand the underlying mechanism, we carried out ChIP-seq for H3K4me3 and H3K27me3. We found that *Ino80* deletion led to dramatic reductions in H3K27me3 occupancy at INO80-bound bivalent gene promoters (Fig. 3d-e, 3h) in the primed state, and we validated such changes by ChIP-qPCR (Fig. 3i). In contrast, the deletion only slightly affected H3K4me3 occupancy (Fig. 3f-g). Together, our data strongly suggested that INO80 is required for the maintenance of H3K27me3 at bivalent promoters for the poised expression of downstream genes in the primed state. Interestingly, in comparison to the primed state, *Ino80* deletion only had very subtle effects on H3K27me3 or H3K4me3 occupancy in the naïve state, (Supplementary Fig. 3a-b), which likely explains the muted gene expression changes (Supplementary Fig. 3c-d, 1g).

During the transition from the naïve to the primed state, the number of bivalent promoters increases significantly (Fig. 4a) and the increase can be mainly attributed to the gradual gain of H3K27me3 (Fig. 4b, and supplementary fig. 3e-f). This is largely similar to what was observed during mouse embryonic development (23,24). We found that the majority of the INO80-bound bivalent genes in the primed state, such as *T*, were already occupied by INO80 in the naïve state (Fig. 4c-e, and supplementary Fig. 3g), suggesting that INO80 may be involved in the establishment of H3K27me3 at these regions. To test this, we carried out H3K27me3 ChIP-seq during the naïve to primed transition in wild-type and *Ino80* deletion cells. We found that Ino80 deletion led to significant cell loss (Fig. 4f). Furthermore, it resulted in impaired H3K27me3 deposition at INO80-bound bivalent promoters (Fig. 4g-i), as well as the de-repression of bivalent genes (Fig. 4j). Compared to wild type cells in which 55% bivalent promoters were established after 2 days of culture in FA, *Ino80* KO cells had only 21% detectable bivalent promoters (Fig. 4k). Within the detected bivalent domains, *Ino80* KO cells also showed much reduced H3K27me3 signal (Fig. 4l). Therefore, INO80 is required not only for the maintenance of H3K27me3 at bivalent promoters in the primed state, but also for the proper establishment of H3K27me3 and bivalency at key developmental genes during the naïve to primed transition.

**Figure 4.**
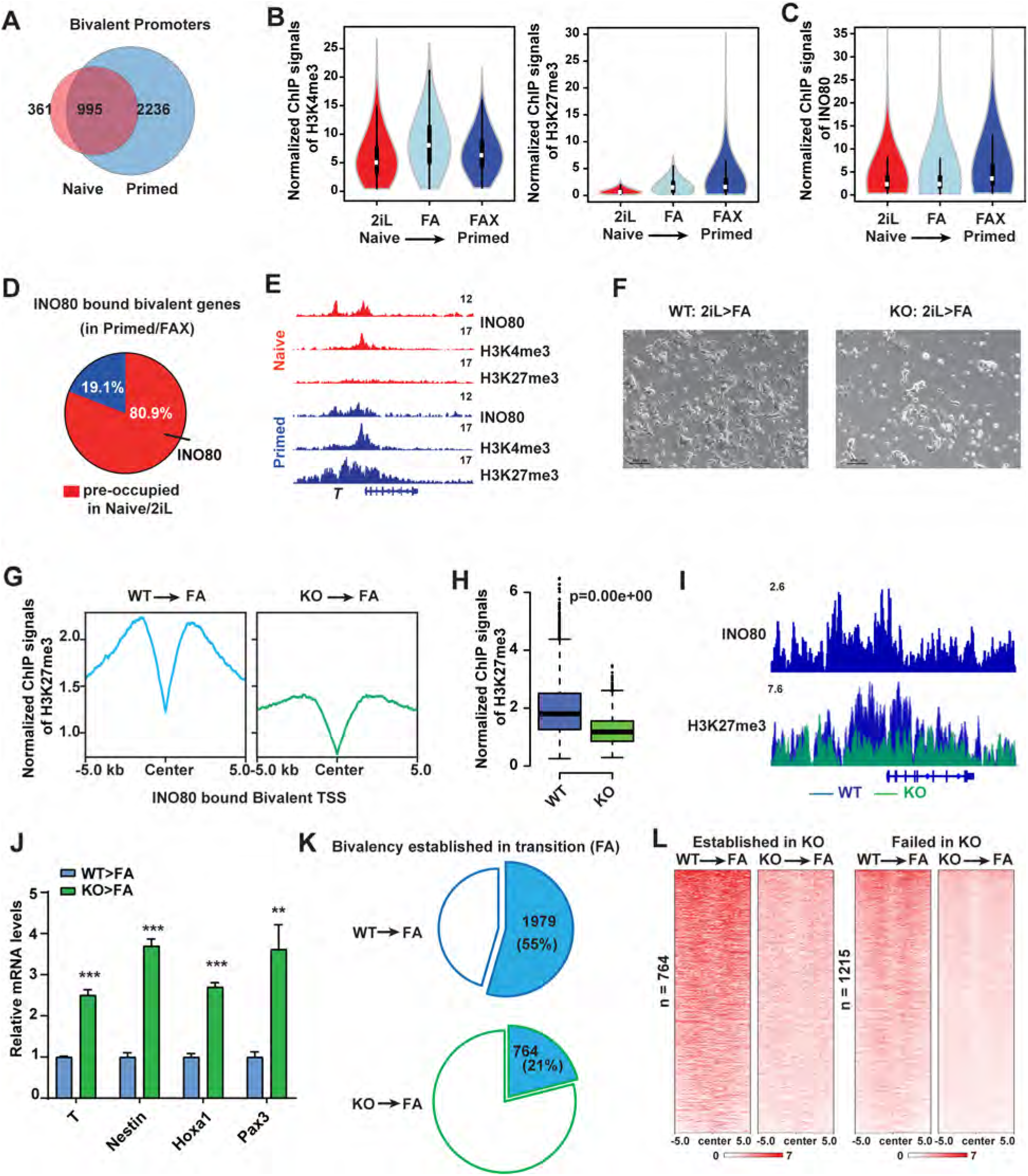
INO80 is required for the establishment of bivalency. **(a)** Venn diagram to show the overlap of bivalent promoters between the naïve and primed state. **(b)** Violin plots to show H3K4me3 and H3K27me3 ChIP-seq signal intensity at bivalent promoters during the naïve to primed transition. **(c)** Violin plots to show INO80 ChIP-seq signal intensity at bivalent promoters during the naïve to primed transition. **(d)** Pie chart to show INO80 occupancy in the naïve state at the primed bivalent promoters. **(e)** Genome browser view of INO80, H3K4me3, H3K27me3 occupancy at T in the naïve and primed state. **(f-l)** *Ino80* deletion during the naïve to primed transition. ESCs were cultured in 2iL and treated with 4-OHT to induce *Ino80* deletion for two days. Cells were then transferred to the FA medium for another 2 days, imaged (f), and collected for ChIP-seq and RT-qPCR (g-i). H3K27me3 occupancy at INO80-bound primed bivalent gene promoters was examined by metagene (g) and box plots (h). p-value was calculated by Wilcoxon signed rank test. INO80 and H3K27me3 occupancy near T was shown by the genome browser view (i). The relative expression of selected primed bivalent genes in WT and *Ino80* deletion cells was determined by RT-qPCR, normalized to *Gapdh* and plotted as mean ± SEM. p-values were calculated by student *t*-test: * <0.05, ** <0.01, *** <0.001. The numbers of successfully established bivalent promoters during the naïve to primed transition in WT or *Ino80* deletion cells were shown by the pie chart (k). H3K27me3 ChIP-seq signal at INO80-bound primed bivalent gene promoters in WT and *Ino80* deletion cells were shown by heatmap and sorted based on signal intensity (l).

### INO80 promotes H2A.Z occupancy as a chromatin remodeler

Next, we wanted to investigate how INO80 regulates H3K27me3 occupancy. Because INO80 is a chromatin remodeler, we hypothesized that it regulates H3K27me3 via its chromatin remodeling function. To test the hypothesis, we introduced dox-inducible wild-type or ATPase-dead INO80 mutants (K551A or E665Q)(39,40) into the *Ino80*-cKO ESCs. As expected, the expression of wild-type INO80 rescued the differentiation and cell death caused by *Ino80* deletion (Fig. 5a). However, the expression of ATPase-dead mutants, albeit at similar levels (Supplementary Fig. 4a), was not able to maintain the cells under the same condition (Fig. 5a). In addition, we also mutated the endogenous *Ino80* gene and generated an INO80 ATPase-dead knock-in ESC line (the K551A/E665Q double mutant, or *Ino80*-KA/ED) by CRISPR-mediated genome editing. The mutations did not drastically affect the INO80 protein level (Supplementary Fig. 4b). However, similar to *Ino80* deletion, *Ino80* ATPase-dead mutations led to cell loss (Supplementary Fig. 4c) and elevated expression of bivalent genes during transition toward the primed state (Supplementary Fig. 4d). Together, the above results indicated that the chromatin remodeling activity of INO80 is essential for the transition toward the primed state and the poised expression of bivalent genes.

**Figure 5.**
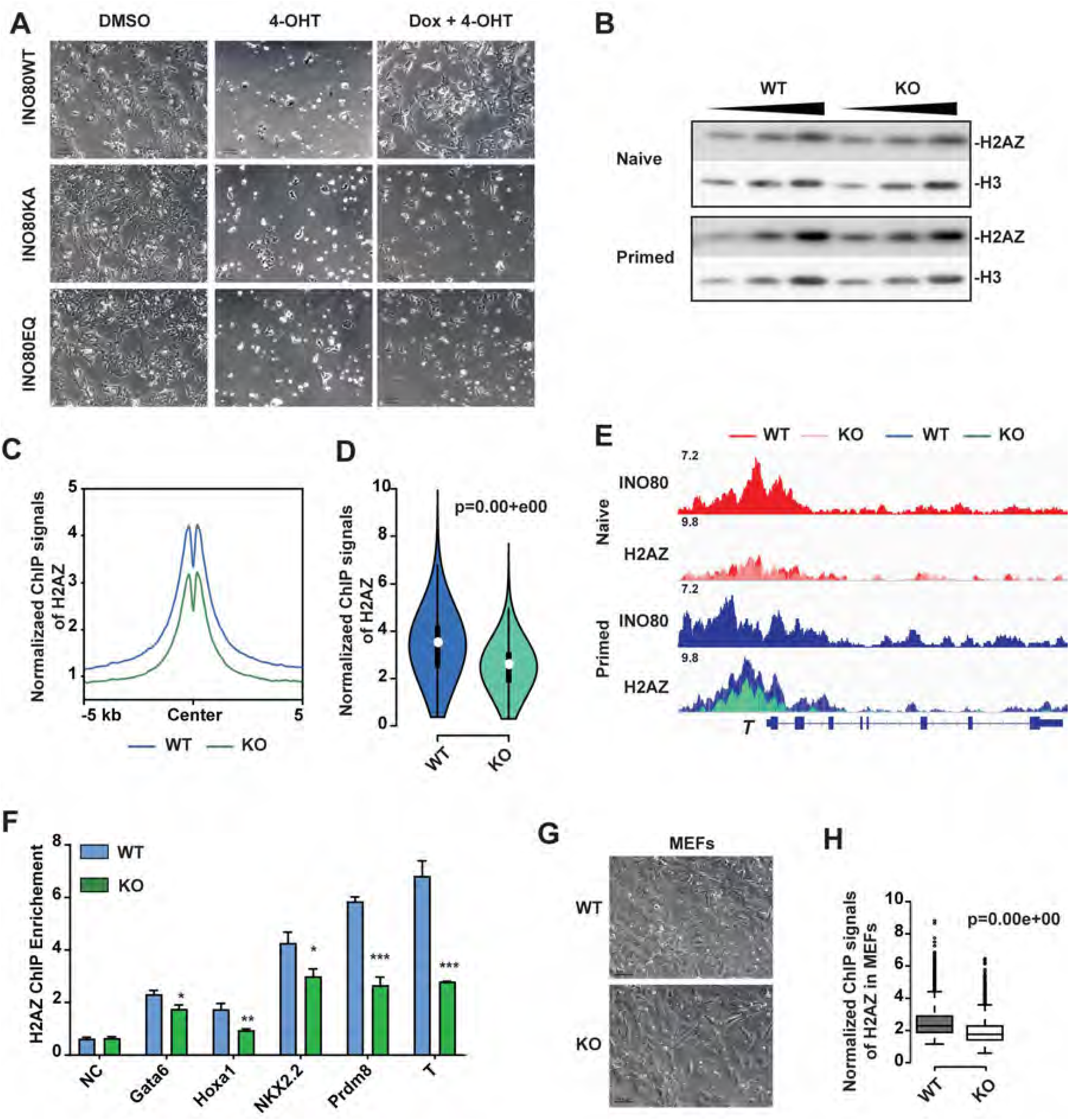
INO80 promotes H2A.Z occupancy. **(a)** Rescue of *Ino80* deletion phenotype by WT INO80 or INO80-ATPase-dead mutants*. Ino80* deletion ESCs were transfected with the piggyBac vectors expressing Dox-inducible WT or ATPase-dead *Ino80* (K551A, KA or E665Q, EQ). *Ino80* deletion was induced by 4-OHT treatment and exogenous *Ino80* expression was induced by Dox treatment simultaneously in serum/Lif. Cell morphology was imaged after one passage. Scale bar = 100 μm. **(b)** Western blots of H2A.Z in WT and Ino80 deletion cells. Histone H3 was used as a loading control. **(c-d)** H2A.Z occupancy in WT and Ino80 deletion cells in the primed state. ESCs were cultured in FAX and treated with DMSO or 4-OHT for 2 days and collected for ChIP-seq. Metagene (c) and Violin plot (d) of normalized H2A.Z ChIP-seq signals at the INO80-bound bivalent promoters. p-values was calculated by Wilcoxon signed rank test. **(e)** ChIP-qPCR to show H2A.Z occupancy at representative bivalent gene promoters in WT and *Ino80* deletion cells. Fold-enrichment was plotted as mean ± SEM. p-values were calculated by student *t*-test: * <0.05, ** <0.01, *** <0.001. **(f)** Genome browser view of INO80 and H2A.Z occupancy near T. **(g-h)** H2A.Z occupancy in WT and Ino80 deletion MEFs. MEFs were treated with DMSO or 4-OHT for 2 days. Cells were imaged (g) and collected for H2A.Z ChIP-seq. Normalized H2A.Z ChIP-seq signals at TSSs was shown in the box plot (h).

As a remodeler, INO80 can regulate nucleosome positioning and composition. To test which of these functions is involved, we first examined the role of INO80 in nucleosome occupancy by MNase titration followed by high throughput sequencing(41) (Supplementary Fig. 4e). To our surprise, *Ino80* deletion only had very limited impact on nucleosome positioning and density (Supplementary Fig. 4f). Moreover, the small changes in nucleosome signals after *Ino80* deletion were similar in both the naïve and primed state and showed no clear correlation with gene expression changes. Thus, changes in the nucleosomes could not explain the *Ino80* deletion phenotypes.

Next, we examined the role of INO80 in the histone variant exchange, as INO80 has been reported to control the eviction of H2A.Z(30,31). Using public ChIP-seq data in mouse ESCs(42), we found that INO80 and H2A.Z showed the highest correlation in genomic occupancy among all the examined chromatin remodelers including BRG1, MBD3, EP400, TIP60, CHD1, CHD2, CHD4, CHD6, CHD7, CHD8 and CHD9 (Supplementary Fig. 5a). This co-localization is consistent with the notion that INO80 may be an important regulator for H2A.Z. We then carried out H2A.Z ChIP-seq in wild-type and *Ino80* deletion cells. In wild-type cells, a sizable portion of H2A.Z localized at TSSs (Supplementary Fig. 5b) and its signal intensity was much higher in the primed state compared to the naïve state (Supplementary Fig. 5c). In *Ino80* deletion cells, although the H2A.Z protein level did not change (Fig. 5b), its occupancy at INO80-bound TSSs was significantly reduced in the primed state (Fig. 5c-e). We validated the reductions by ChIP-qPCR at promoters of representative bivalent genes (Fig. 5f). Therefore, INO80 is required for the maintenance of H2A.Z in the primed state. Interestingly, reminiscent of what was observed for H3K27me3 (Supplementary Fig. 3a), *Ino80* deletion had minimal impact on H2A.Z occupancy in the naïve state (Supplementary Fig. 5d-e), possibly due to the already low basal H2A.Z occupancy (Supplementary Fig. 5c). This result provided a possible explanation for why Ino80 is dispensable for H3K27me3 binding in the naïve state.

Because our results showed an opposite role of INO80 in H2A.Z occupancy compared to earlier studies(30,31), we wondered whether our observation was specific to pluripotent stem cells. We thus tested the impact of *Ino80* deletion in mouse embryonic fibroblasts (MEFs, Fig. 5g). Similar to the above results, *Ino80* deletion led to significant reduction in H2A.Z signals in MEFs as well (Fig. 5h). Therefore, the requirement for INO80 in H2A.Z genomic occupancy appears to be conserved among different cellular states in mammalian cells.

### INO80-dependent H2A.Z occupancy licenses the establishment of bivalency

Based on the above data, we hypothesized that INO80-dependent H2A.Z occupancy is a determining factor in bivalent promoter regulation. Consistent the notion, INO80 pre-occupied 80.9% of all bivalent promoters in the primed state (Fig. 4d). Moreover, almost all the INO80-bound bivalent promoters were co-occupied by H2A.Z (Supplementary Fig. 6a) and showed stronger H2A.Z signal (Supplementary Fig. 6b). To test the hypothesis, we generated H2A.Z conditional deletion ESCs, in which the isoform *H2az2* was constitutively deleted and *H2az1* could be deleted upon 4-OHT treatment (Supplementary Fig. 6c-e). Similar to *Ino80* deletion, *H2az1/H2az2* double deletion had limited impact on the naïve state but led to severe cellular differentiation and cell loss in the transition to primed state (Fig. 6a). Importantly, in the primed state, the double deletion resulted in impaired H3K27me3 occupancy at bivalent promoters (Fig. 6b-d) and the impairment was more obvious at regions with stronger H3K27me3 signal (Supplementary Fig. 6f). Consistently, *H2az1/H2az2* deletion caused de-repression of bivalent genes in the primed state (Fig. 6e), as well as premature activation of developmental genes during subsequent embryoid body differentiation (Fig. 6f-g), phenocopying *Ino80* deletion (Fig. 1f and 3c). Therefore, H2A.Z is required for the maintenance of H3K27me3 and the poised expression of bivalent genes.

**Figure 6.**
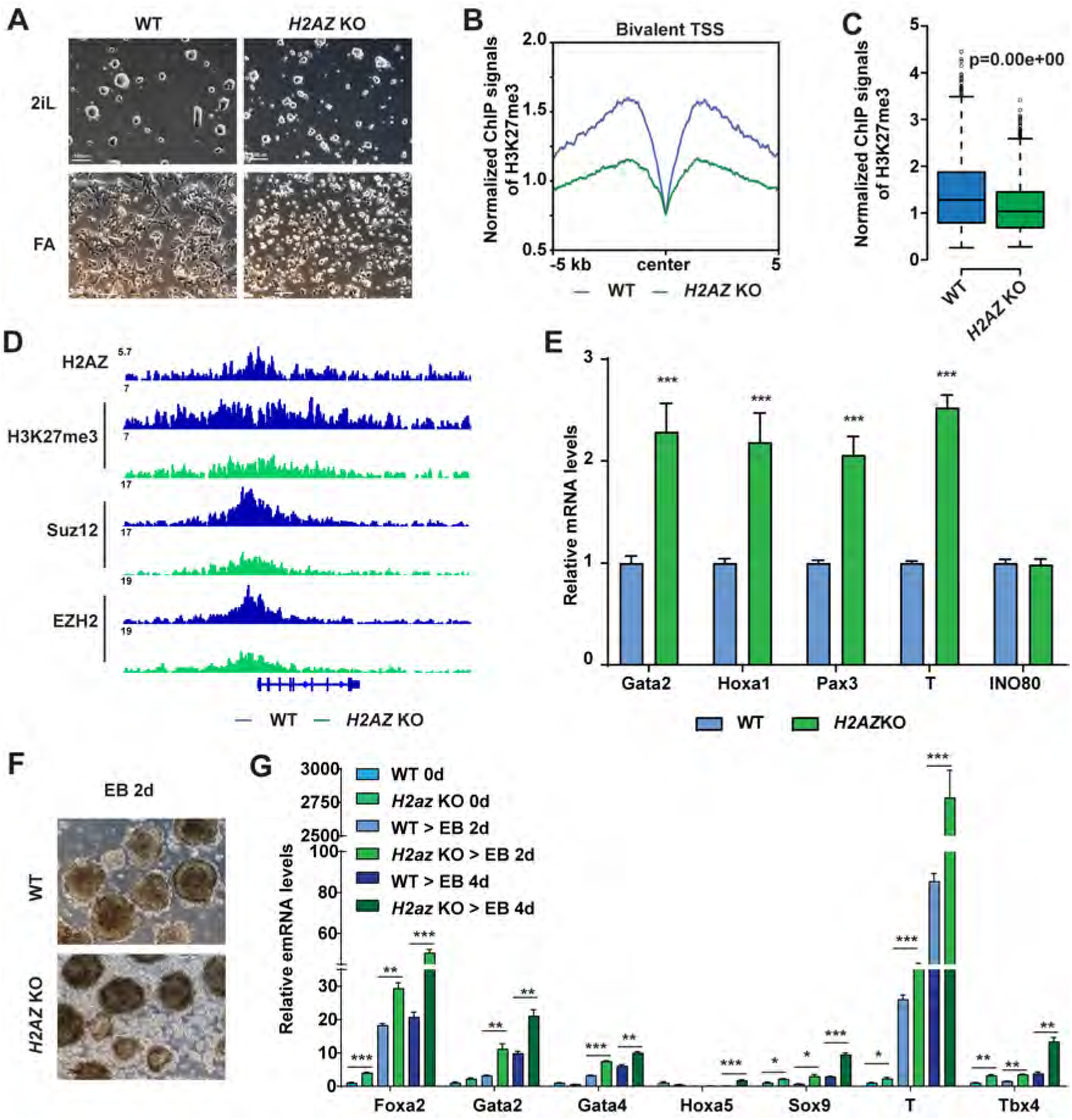
INO80-dependent H2A.Z deposition licenses the establishment of bivalency. **(a)** *H2az1/H2az2* deletion in the naïve state and during the naïve to primed transition. *H2az1*-cKO/*H2az2*-KO ESCs were cultured in the naïve state in 2iL and treated with DMSO or 4-OHT for 2 days. Cells were then maintained in the naïve state in 2iL or transitioned into FA for another 2 days to initiate the primed state. Cells were imaged and collected for ChIP-seq and RT-qPCR experiments. Scale bars = 100 μm. **(b-c)** H3K27me3 occupancy at INO80-bound bivalent TSSs in WT and *H2az1/H2az2* deletion cells. Metagene and box plots were generated using normalized H3K27me3 ChIP-seq signals. p-value was calculated by Wilcoxon signed rank test. **(d)** Genome browser view of H2A.Z, H3K4me3, H3K27me3, PRC2 component SUZ12 and EZH2 near T. **(e)** Expression of representative bivalent genes in WT and *H2az1/H2az2* deletion cells. Relative mRNA levels were determined by RT-qPCR, normalized to *Gapdh* and plotted as mean ± SEM. p-values were calculated by student *t*-test: * <0.05, ** <0.01, *** <0.001. **(f-g)** Expression of representative bivalent genes in WT and *H2az1/H2az2* deletion cells during EB formation. WT or *H2az1*/*H2az2* deletion cells were induced to transition toward the primed state as described in A. Cells were aggregated to form EB for 2 days, imaged (f), and collected for RT-qPCR. Scale bars = 100 μm. Expression of representative bivalent genes in EB differentiation was determined by RT-qPCR, normalized to *Gapdh* and plotted as mean ± SEM (g). p-values were calculated by student *t*-test: * <0.05, ** <0.01, *** <0.001.

To understand how INO80 and H2A.Z regulates H3K27me3, we carried out ChIP-seq for key components of the PRC2 complex in wild-type, *Ino80* deletion, and *H2az1/H2az2* deletion cells in the primed state. We found that both *Ino80* and *H2az1/H2az2* deletion impaired the chromatin binding of the PRC2 structural component SUZ12 (Fig. 7a-e). In addition, *H2az1/H2az2* deletion also impaired the binding of the PRC2 enzymatic subunit EZH2 (Fig. 7f-g). We validated these results by ChIP-qPCR (Fig. 7h). In contrast, H2A.Z occupancy was not significantly affected by the deletion of PRC2 components EZH2 or EED (Supplementary Fig. 7a-c). These results suggested that INO80 and H2A.Z are required for the recruitment of a fully-assembled PRC2 for H3K27me3 deposition at bivalent promoters. To add further support, we analyzed the genomic occupancy by chromatin factors in ESCs using public data(42,43). We found that INO80 closely clustered with H2A.Z and PRC2 at bivalent promoters among major chromatin remodelers (Fig. 7i). Besides INO80, the NuRD complex has been shown to repress bivalent genes via H3K27ac deacetylation(44). As expected, the core NuRD components CHD4 and Mbd3 also showed strong colocalization with PRC2, supporting the validity of this analysis.

**Figure 7.**
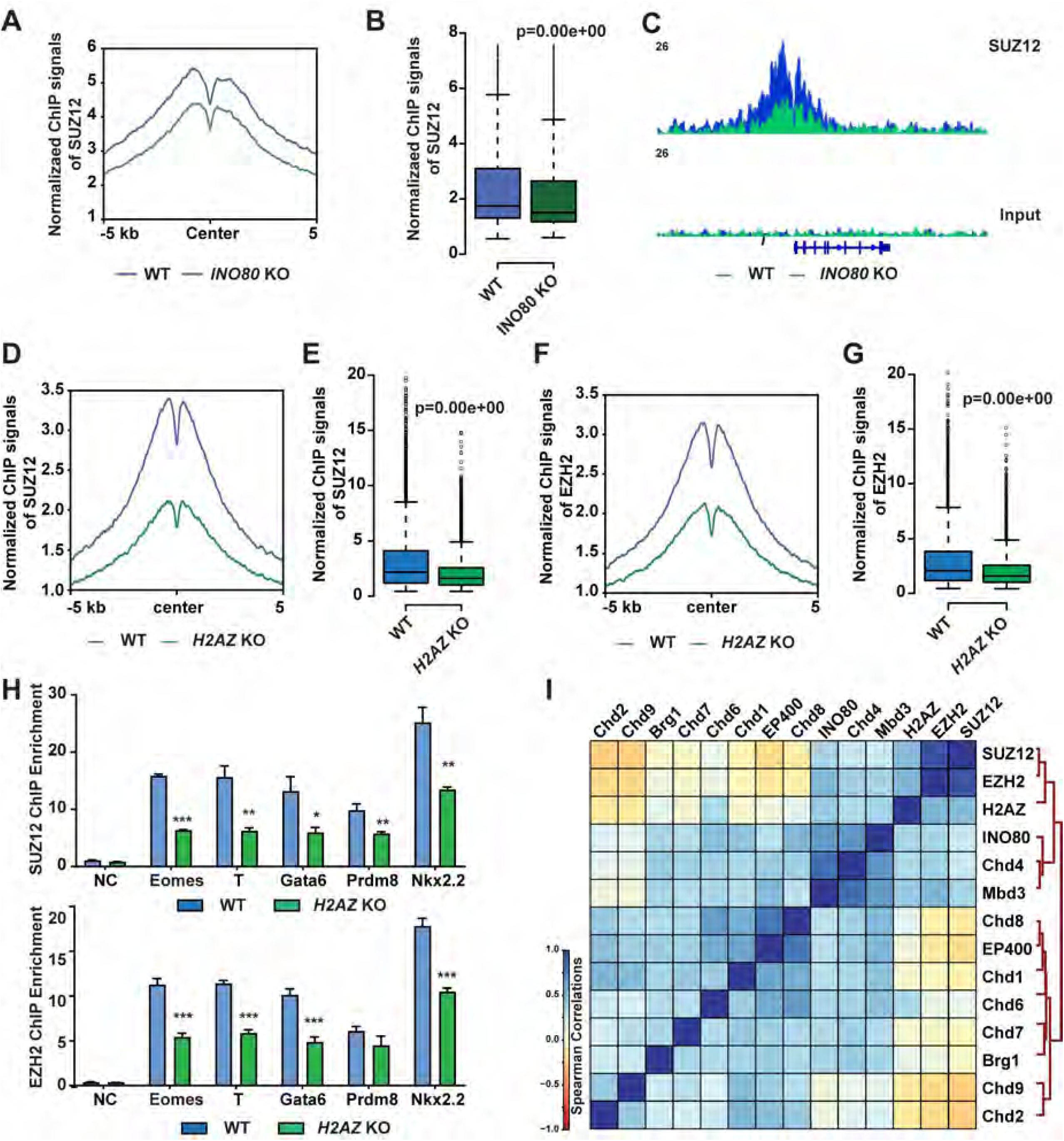
INO80-dependent H2A.Z occupancy is required for PRC2 recruitment. **(a-b)** SUZ12 occupancy at INO80-bound bivalent TSSs in the primed state in WT and *Ino80* deletion cells. *Ino80*-cKO ESCs were cultured in FAX and treated with 4-OHT for 2 days. Cells were cultured for 2 additional days and collected for ChIP-seq. SUZ12 occupancy was shown by metagene (a) and box plots (b) using normalized ChIP-seq signals. **(c)** Genome browser track to show SUZ12 occupancy near T in WT and *Ino80* deletion cells. **(d-g)** SUZ12 and EZH2 occupancy at INO80-bound bivalent TSSs in the primed state in WT and *H2az1/H2az2* deletion cells. *H2az1*-cKO/*H2az2*-KO ESCs were cultured in the primed state in FAX and treated with DMSO or 4-OHT for 2 days. Cells were cultured for another 2 days and collected for ChIP-seq. SUZ12 and EZH2 occupancy was shown by metagene (d, f) and box plots (e, g) using normalized ChIP-seq signals. p-value was calculated by Wilcoxon signed rank test. (**h**) ChIP-qPCR to show SUZ12 (upper) and EZH2 (lower) at representative bivalent gene promoters in WT and *Ino80* deletion cells. Fold enrichment was plotted as mean ± SEM and p-values were calculated by student *t*-test: * <0.05, ** <0.01, *** <0.001. **(i)** Spearman correlation of genomic occupancy among chromatin factors. Except for that of INO80 and H2AZ, genomic occupancy of all other chromatin factors was based on public datasets.

Taken together, we propose the following model: During the naïve to primed transition, INO80 occupancy at TSSs near developmental genes promotes the deposition of H2A.Z. In turn, increased H2A.Z binding facilitates the recruitment of PRC2 and methylation of H3K27 to establish bivalent domains. The bivalent histone modifications maintain the developmental genes in a poised state for activation during further differentiation and development. Our model strongly argues that INO80-dependent H2A.Z deposition is a critical licensing step in bivalent chromatin establishment and maintenance during cell fate transitions.

## Discussion

ATP-dependent chromatin remodeling complexes play important roles in pluripotent stem cells. While many interact with and target specific nucleosomes(42), we showed that INO80 does not appear to drastically change global nucleosome positioning or density. Instead, it promotes H2A.Z occupancy and licenses the formation and maintenance of bivalent domains at developmental gene promoters. Our findings support the notion that different remodelers use different mechanisms and act on different genes to maintain the transcription program in pluripotent stem cells.

Our findings are also consistent with the mouse phenotypes. Specifically, *Ino80* deletion led to post-implantation lethality and the embryos failed to develop beyond the egg cylinder stage(32,35). Deletion of H2A.Z and PRC2 components resulted in developmental arrest around gastrulation(45–47). The similarities in the developmental defects and timing in the animal models support the idea that INO80, H2A.Z and PRC2 coordinately regulate gene expression in post-implantation embryonic development. They also support our result that *Ino80* is required in the primed pluripotent state and bivalent gene regulation. Interestingly, *Ino80* is dispensable for the naïve pluripotent state, but is necessary to maintain ESC self-renewal in the serum/LIF condition(33). The differences may likely lie in the specific cellular state, as ESCs cultured in serum/LIF are heterogenous and known to require many more factors that are otherwise expendable in the naïve state(3,48). In contrast, the naïve state is thought to be the ground state that is more robust and stable(49). It is characterized with an epigenome that has low levels of DNA and repressive histone methylations(5,6), and it can be maintained without many of the repressive epigenetic mechanisms(50). Consistently, we found that H2A.Z genomic occupancy is low in the naïve state and *Ino80* deletion did not further reduce its level significantly, which may explain why *Ino80* is not required under this condition.

INO80 was initially thought to facilitate H2A.Z eviction(30,31). However, other studies in yeast did not detect large changes in H2A.Z binding upon Ino80 deletion(51,52). In mouse ESCs, we found that INO80 showed the highest co-localization with H2A.Z in the genome among major ATP-dependent chromatin remodelers. Furthermore, INO80 pre-marks genomic regions in the naïve state before significant levels of H2A.Z is established, and it is required for the maintenance of H2A.Z occupancy in the primed state. Finally, *Ino80* deletion led to reduced H2A.Z occupancy in MEFs as well, suggesting that its role in H2A.Z deposition and maintenance is not unique to ESCs. Consistent with our data, a recent study showed that *Ino80* mutant caused decreased H2A.Z binding at key flowering genes in Arabidopsis(53). Thus, INO80 likely plays a complex role in H2A.Z regulation especially in higher eukaryotes.

The histone variant H2A.Z can activate or repress downstream genes in a context dependent manner and is involved in many developmental and disease processes(54). In serum/LIF cultured ESCs, H2A.Z is enriched at gene promoters and PRC2 target genes, and it facilitates both positive and negative regulatory complexes to access chromatin and regulate genes important for ESC self-renewal and differentiation(55–57). Consistently, we showed that H2A.Z binds both active and repressed gene promoters in the naïve and primed pluripotent state. However, we found that H2A.Z shows stronger signals at the bivalent gene promoters, and it is selectively required in the primed but not the naïve state. This is consistent with the phenotype of H2A.Z deletion in mice, in which the null embryos were normal up to E4.5 and started to show defects after implantation(45). Moreover, we identified H2A.Z as crucial prerequisite in H3K27me3 deposition and bivalent domain formation. Our results provided additional support to earlier reports on the interplay between H2A.Z and H3K27me3(55,57,58). More importantly, they provided a possible mechanism to explain the *de novo* formation of bivalent domains after H3K27me3 erasure(22), in that H2A.Z may serve as a bookmark in the absence of H3K27me3. Finally, we showed that H2A.Z occupancy is low in the naïve state, and we think it possibly contributes to the low H3K27me3 modification at bivalent promoters despite high expression of PRC2 components(11).

The bivalent modifications by H3K4me3 and H3K27me3 at developmental gene promoters are thought to keep downstream genes in a poised state(19,20). In agreement with this notion, we showed that reduced H3K27me3 occupancy at bivalent promoters in *Ino80* or *H2az1/H2az2* deletion cells led to de-repression of the downstream genes and disruption of the primed pluripotent state. Our data supports the functional significance of the bivalent domains in maintaining an intricate balance between self-renewal and differentiation in pluripotency and early development. Indeed, recent reports showed that bivalent modifications are largely absent from developmental gene promoters in mouse pre-implantation embryos but become enriched soon after implantation, and a strong bivalent chromatin is a hallmark for the primed pluripotent state(23,24). In summary, we identified INO80 as an essential regulator for bivalent domains and primed pluripotency via H2A.Z. Because INO80 and H2A.Z have been heavily implicated in development and disease, it is conceivable that the INO80-H2A.Z-H3K27me3 axis may play important roles in a similar fashion during other cell fate transitions.

## Methods

### Cell Culture

For routine culture, mouse ESCs were maintained in the ESGRO medium (Millipore) on tissue culture plates pretreated with 0.1% gelatin. For the naïve pluripotent state, mouse ESCs were cultured in 2iL medium containing N2B27 medium supplemented with PD0325901 (1 mM, Selleck), CHIR99021 (3 mM, Selleck) and 1000 units/ml recombinant murine LIF (Millipore) on gelatin-coated plates (2iL). For the primed pluripotent state, mouse ESCs were cultured in FA or FAX medium containing N2B27 supplemented with FGF2 (Peprotech, 12 ng/ml), Activin A (Peprotech, 20 ng/mL), and XAV-939 (2 uM Selleck) on human fibronectin-coated plates (Millipore). The N2B27 medium contains (1000 ml): 500 ml DMEM/F12 (Invitrogen; 11320), 500 ml Neurobasal (Invitrogen; 21103), 5 ml N2 supplement (Invitrogen; 17502048), 10 ml B27 supplement (Invitrogen; 17504044) and 0.1% bovine serum albumin fraction V (Thermofisher).

### Mouse embryonic stem cell lines

The E14Tg2a ESCs were obtained from MMRRC. The *Ino80* conditional deletion ESC line (Ino80^flox/flox^;UBC-CreERT2) was derived from E3.5 blastocysts from the breeding of Ino80^flox/+^;UBC-CreErt2 and Ino80^flox/+^ in the 129S1/SvImJ background. The Ino80^flox/flox^ mouse strain was obtained from blastocyst injection of the Ino80^flox/flox^ ESCs ordered from EuMMCR. All animal experiments were approved by the NIEHS Institutional Animal Care and Use Committee and were performed according to the NRC Guide for the Care and Use of Laboratory Animals. ESC liness expressing the INO80 ATPase-dead mutants were generated by transfecting the *Ino80* conditional deletion ESCs with piggyBac vectors (kindly provided by Dr. Joanna Wysocka, Stanford University) containing the corresponding Ino80 mutant cDNAs and the PB transposase (System Biosciences). The INO80-HA and INO80 ATPase-dead knock-in ESCs were generated by CRISPR-mediated genome editing. Briefly, sgRNAs targeting the desired genomic loci were cloned into pX330 (addgene #42230). Single-stranded oligo DNA donor (ssODN) homologous recombination templates were synthesized by IDT. sgRNAs and ssODN templates were co-transfected into E14Tg2a cells, and correctly targeted clones were identified by PCR genotyping and confirmed with secondary assays. The *H2az1* flox/flox; *H2az2 Δ/Δ;* Rosa26-CreERT2 (H2az1-cKO/H2az2-KO) ESC line was generated by CRISPR-mediated genome targeting in the Rosa26:CreERT2 ESCs (kindly provided by Dr. Shaun Cowley, University of Leicester). To induce gene deletion, 100 nM 4-OHT (Selleck) was added to culture medium for two days.

### Embryoid Body formation assay

For embryoid body (EB) formation, ESCs were induced to primed state in FA for two days. Simultaneously, cells were treated with DMSO or 0.1 mm 4-OHT. Then 500,000 cells/ml single cell suspension in DMEM medium containing with 10% FBS were added to ultra-low attachment plate (Corning) in a 37°C, 5% CO2 incubator to allow them to form EBs. Medium were refreshed every other day. EBs were collected at indicated days.

### Protein extract and Western blotting

Cells were lysed in RIPA buffer (ThermoFisher) containing protease inhibitor cocktail (PIC, Roche) and PMSF, and total protein concentration was determined using the BCA kit (Pierce). Cell lysate was loaded into a NuPAGE^®^ Bis-Tris gels and transferred onto nitrocellulose membrane. The blot was blocked with 5% non-fat milk at room temperature for 30 min, followed by incubation with primary antibodies at 4°C overnight. The blot was subsequently incubated with either horse-radish peroxidase (HRP)-conjugated antimouse or rabbit IgG. Signal was detected using ChemiDoc Imaging system (Bio-Rad).Antibodies against specific antigens are provided in Supplementary Table 1. All experiments were repeated 2-3 times and one representative result was shown in the figures.

### RNA isolation, quantitative real-time PCR and RNA-seq

Total RNA was extracted using the GeneJET RNA purification kit (ThermoScientific). RNA was reverse transcribed using iScript cDNA Synthesis Kit (Bio-Rad). Quantitative real-time PCR (qPCR) were performed with the SsoAdvanced Universal SYBR Green Supermix (Bio-Rad) on the CFX384 real-time PCR detection system. All experiments were carried out with at least two biological replicates. For RNA-seq with spike-in controls, 10 ul of 1:100 dilution of the ERCC Spike-In Mix #1 was added to the lysate of 1 million cells before RNA extraction. Total RNA was then extracted as described above. For RNA-seq library preparation, 1 mg total RNA was used to prepare the RNA-seq library with the Truseq RNA Library Prep Kit V2 with Ribo-Zero (Illumina). Primers used in this study are provided in Supplementary Table 2. All experiments were repeated 3 times and one representative result was shown in the figures.

### ATAC-seq, MNase ChIP-seq, MNase-seq and ChIP-qPCR

Omni-ATAC was performed with Nextera DNA library prep kit (Illumina) as described(59). Then tagmentated DNA was purified using DNA Clean & Concentrator-5 Kit (Zymo). ATAC-seq library was generated by PCR amplification and purified with AMPure XP beads (Beckman Coulter).

For ChIP-seq, mouse ESCs were digested by trypsin (Naive state) or Accutase (Primed state) to single cells and fixed in 1% formaldehyde for 10 min at room temperature. Crosslinking was quenched by the addition of glycine to the final concentration of 0.25M and the incubation at room temperature for 5 min. Cells were washed twice with ice-cold PBS containing the protease inhibitor cocktail (PCI, Roche). Cell pellet was resuspended and permeabilized with hypotonic buffer (10 mM HEPES, 10 mM KCl, 1.5 mM MgCl2, 0.34 M sucrose, 10% glycerol, 1% Triton X-100, and freshly add 1 mM DTT, 0.5% N-P40 and Protease inhibitors) on ice for 10 min. Cells were pelleted by centrifugation at 1350 x g, washed once with the hypotonic buffer, and again with the MNase buffer (20 mM Tris-HCl pH 7.5, Sucrose 0.34 M, KCl 60 mM, NaCl 15 mM and 1 mM CaCl2). Cell pellet was resuspended in the MNase buffer and digested with MNase (100-200 units per 10 M cells) at 37 °C for 12 min on a shaker. Digestion was stopped by the addition of 10% volume of 0.5M EDTA and mixed with one volume of 2X lysis buffer (20 mM Tris-HCl pH8.0, 300 mM NaCl, 2 mM EDTA, 20% Glycerol, 1% Triton X-100, 2X PIC, 0.2mM PMSF). Cells were sonicated by 4 cycles of 30 s ON and 90 s OFF on the Misonix S3000 sonicator with 30-watt power output, and insoluble materials were removed by centrifugation at 21000 x *g* at 4°C for 20 min. 25 ug chromatin was used for each ChIP experiment, and was incubated with the primary antibody (Supplementary Table 2) and the protein A/G Dynabeads (ThermoFisher) overnight at 4°C. The beads were washed with low salt buffer (20 mM Tris–Cl (pH 8.0), 150 mM NaCl, 1 mM EDTA, 1% Triton X-100), high salt buffer (20 mM Tris– Cl (pH 8.0), 250 mM LiCl, 1 mM EDTA, 1% Triton X-100) and Lithium Chloride buffer (20 mM Tris–Cl (pH 8.0), 500 mM NaCl, 1 mM EDTA, 1% Triton X-100 and 1% Sodium Deoxycholate), and then washed twice with the TE buffer (10 mM Tris-HCl, 1mM EDTA). Immunoprecipitated DNA-protein complex was eluted with the elution buffer (50 mM Tris–Cl (pH 7.5), 10 mM EDTA, 1% sodium dodecyl sulphate) at 65°C for 15 min. The elution was collected and reverse-crosslinked at 65 °C overnight, diluted with 1 volume of TE buffer and treated with RNase A at 37 °C for 30 min followed by 0.2 μg/ml Proteinase K at 55 °C for 60 min. The immunoprecipitated DNA was purified from the elution by the Zymo DNA Clean & Concentrator Kit. For ChIP-seq, 2 ng ChIP DNA or input was used to generate the sequencing library using the Nextflex Rapid DNA-seq Kit (PerkinEImer) or Nextflex DNA and the resulting libraries were sequenced on the Illumina platform. Two biological replicates were carried out for each experiment, and combined reads were used for further analyses.

For MNase-seq, fixed cells were treated as described above for titrated MNase digestion. Either 4, 16, 64 or 256 U of MNase (NEB) were added to per 2M pre-warmed cells and incubated at 37 °C for 10 min. Digestion was halted by addition of MNase stop buffer (20mM EDTA, 20mM EGTA, 0.4% SDS and 0.5mg/ml Proteinase K). The mixtures were incubated at 65 °C for 60 min to reverse crosslinks. DNA was cleaned up by Gel Purification Column (Qiagen). Fragment size were estimated by Bioanalyzer 2000 (Agilent). 100 ng DNA from each digestion were used to prepare DNA libraries as described in the Nextflex Rapid DNA-seq Kit (PerkinEImer). Size selection was performed after adaptor ligation with Agencourt AMPure XP beads (Beckman Coulter). 8 cycles of PCR were used to amplify the products.

For ChIP-qPCR, either 0.1 ng ChIP DNA or 0.1 ng Input DNA was used as template, and fold-enrichments were determined by the 2^-ΔCT^ method. Primers used in this study are provided in Supplementary Table 2. All experiments were repeated 3 times and one representative result was shown in the figures.

### Data processing and analyses

For all sequencing runs, only reads with Phred quality score >= 20 and length > 15 were kept for analysis. For RNA-seq, reads were first trimmed with cutadapt (v1.9), and filtered reads were aligned to mm9 reference with STAR (v2.5.2b) with parameters ‘--outFilterMismatchNoverLmax 0.04’. Mapped reads were annotated to gene exons to generate a count matrix on gene level with featureCounts (v1.5.1). Differential expressed genes (DEGs) were identified with the R package DESeq2, and DEGs were selected as ļlog2FCļ > 1 and adjusted p value < 0.05. For RNA-seq with ERCC spike-ins, the ERCC reads were aligned separately to the ERCC92 reference. Gene count matrix was generated with the same pipeline as standard RNA-seq. To adjust the RNA-seq result using the ERCC, estimateSizeFactors function in DESeq2 was used with the control genes set to ERCCs. To normalize RPKM of genes to the ERCC spike-in, loess normalization function from R affy package was used to adjust the values. RPKM data for genes were log2 transformed (after adding 1). Gene ontology enrichment of was analyzed by GO Consortium powered by PANTHER.

For ChIP-seq analysis, reads were aligned to mm9 reference using bowtie (v1.1.2) with the following parameters ‘-v 2 −m 1 --best --strata −I 15 −X 1000’ after filtering as described above. PCR duplicates were removed after alignment. For peak calling, SICER (v1.1) was used with window size 200 and gap 600. Peaks were cleaned-up by FDR < 0.0001 and excluded signals from the Encode mm9 blacklist for further analyses. Two-fold enrichment against input was applied to high confidence H3K27me3 peaks. For visualization, Bedtools (v2.21.0) genomeCoverageBed and UCSC utility bedGraphToBigWig were used to build the tracks of the samples. The coverage of the samples was then normalized to the total reads per 10 million for between sample comparison. Deeptools was used for spearman correlation, heatmap and metagene (density) plot. Peaks were annotated to nearest TSS with Homer. Promoter was defined as regions of 500 bp around TSS. Bivalent promoters were defined by promoters with overlapped H3K4me3 and H3K27me3 peaks.

For MNase-seq, reads were aligned using bowtie (v1.1.2) with the same parameters as in ChIP-seq. The reads fragments between 100 to 200 bp were selected for further analyses. For visualization of the nucleosome position pattern in metagene profiles, mid-point of each fragment was selected and followed by median normalization(60). Gaussian smooth was used to smooth the curves in the final plot(61).

For ATAC-seq, same filtering pipeline was used as described in ChIP-seq. The first 9 bp from the end of the aligned fragments were counted as open chromatin reads and normalized to library depth. Reads from chrM were removed.

Software and algorithms were listed in Supplementary Table 3.

### Statistical analysis

RT-qPCR and ChIP-qPCR were analyzed with student *t*-test. RNA-seq, ATAC-seq or ChIP-seq signals were analyzed with Wilcoxon rank sum test (when unpaired) or Wilcoxon signed rank test (when paired).

### Data availability

Genomic data generated for this study have been deposited in the GEO repository with the accession number GSE158545. We used the following published datasets for analysis: CHIP-seq: ATPase chromatin remodelers (GSE64825), SUZ12 and EZH2 (GSE49431), H3K4me3 and H3K27me3 (GSE117896); scRNA-seq: E-MTAB-6967.

## Acknowledgments

We thank Dr. Michael Stadler and Dr. Dirk Schübeler (FMI science) for the help with MNase-seq data normalization. We thank Dr. Joanna Wysocka (Stanford University) for the PiggyBac expression vectors and Dr. Shaun Cowley (University of Leicester) for the Rosa26-CreERT2 ESCs. We thank the NIEHS Epigenomics and DNA Sequencing core, Imaging, and Animal facilities for assistance with various experiments. This study was supported in part by the Intramural Research Program of the NIH, National Institute of Environmental Health Sciences Z01ES102745 (to G.H.).

## Author Contributions

H.Y. and G.H. conceived the study. H.Y., B.L. and G.H. carried out the experiments. J.W., H.Y. and B.B. analyzed the data. H.Y., J.W. B.L., B.B., J.L. and G.H. interpreted the data. H.Y. and G.H. wrote the manuscript.

## Supplementary Figures and figure legends

**Supplementary Figure 1.**
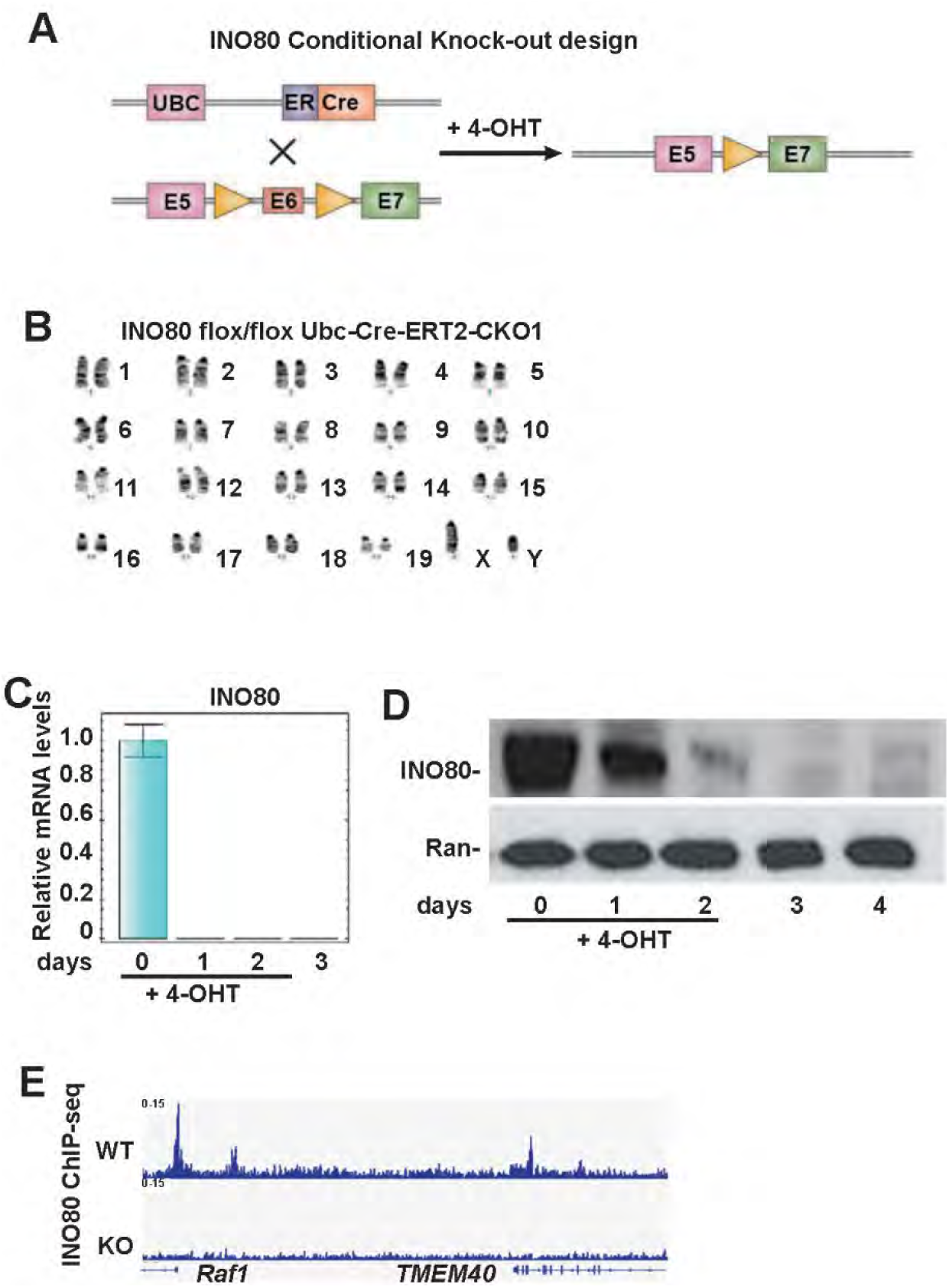
*Ino80* deletion ESCs (related to Figure 1). **(A)** *Ino80* deletion strategy. **(B)** Karyotyping of *Ino80*-cKO ESCs. **(C-D)** *Ino80* expression at the indicated time points after 4-OHT treatment. *Ino80* mRNA expression was determined by RT-qPCR, normalized to *Gapdh*, and plotted as mean ± SEM (C). *Ino80* protein expression was determined by western blot (D). Ran was used as a loading control. **(E)** Genome browser track to show INO80 ChIP-seq in WT and INO80 deletion cells. ESCs were treated with 4-OHT for 2 days, cultured for another 2 days, and collected for ChIP-seq.

**Supplementary Figure 2.**
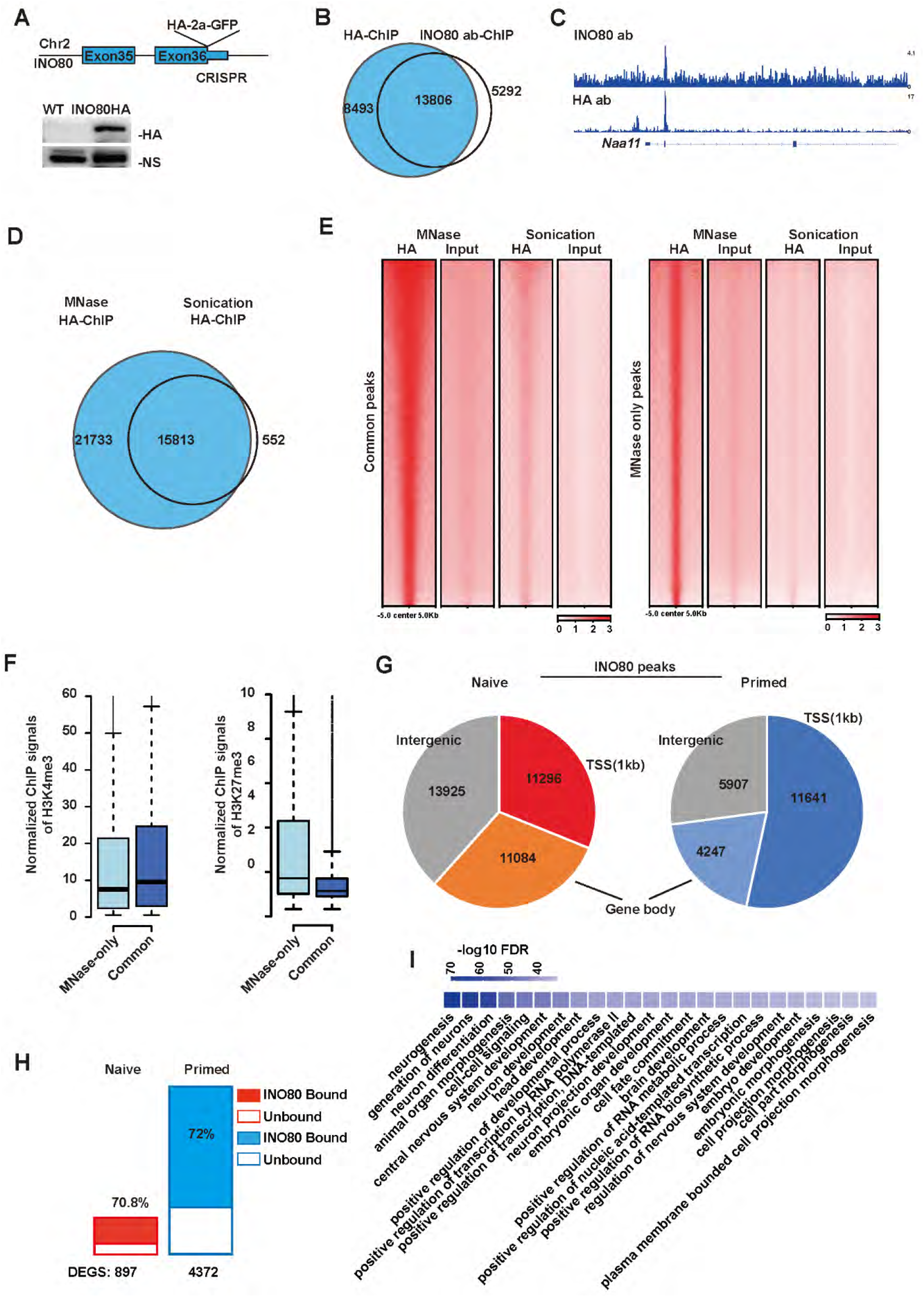
INO80 ChIP-seq (related to Figure 2) **(A)** HA knock-in at the endogenous *Ino80* locus. Upper: targeting strategy; Lower: Western blot using the HA-antibody. NS: a non-specific band from the HA-antibody western was used as the loading control. **(B-C)** Comparison of ChIP-seq using the HA and INO80 antibody in the *Ino80*-HA ESCs. Cells were cultured in serum/LIF. (B) Venn diagram to show the overlap of ChIP-seq peaks. (C) Genome browser track view of ChIP-seq signals. **(D-F)** Comparison of INO80 ChIP-seq by MNase digestion or sonication. (D) Venn diagram to show the overlap of ChIP-seq peaks. (E) Heatmap to show ChIP-seq signals in MNase-ChIP and Sonication-ChIP shared peaks(left) or MNase-specific peaks (right). Peaks were sorted by MNase-ChIP-seq signal intensity. (F) Box plot to show H3K4me3 and H3K27me3 ChIP-seq signals at MNase-specific peaks or MNase-ChIP and Sonication-ChIP shared peaks. **(G)** Distribution of INO80 ChIP-seq peaks in the genome in the naïve and primed state. **(H)** INO80 occupancy near of *Ino80* deletion-induced DEGs in the naïve and primed state. **(I)** Top gene ontology terms enriched in INO80-bound bivalent genes.

**Supplementary Figure 3.**
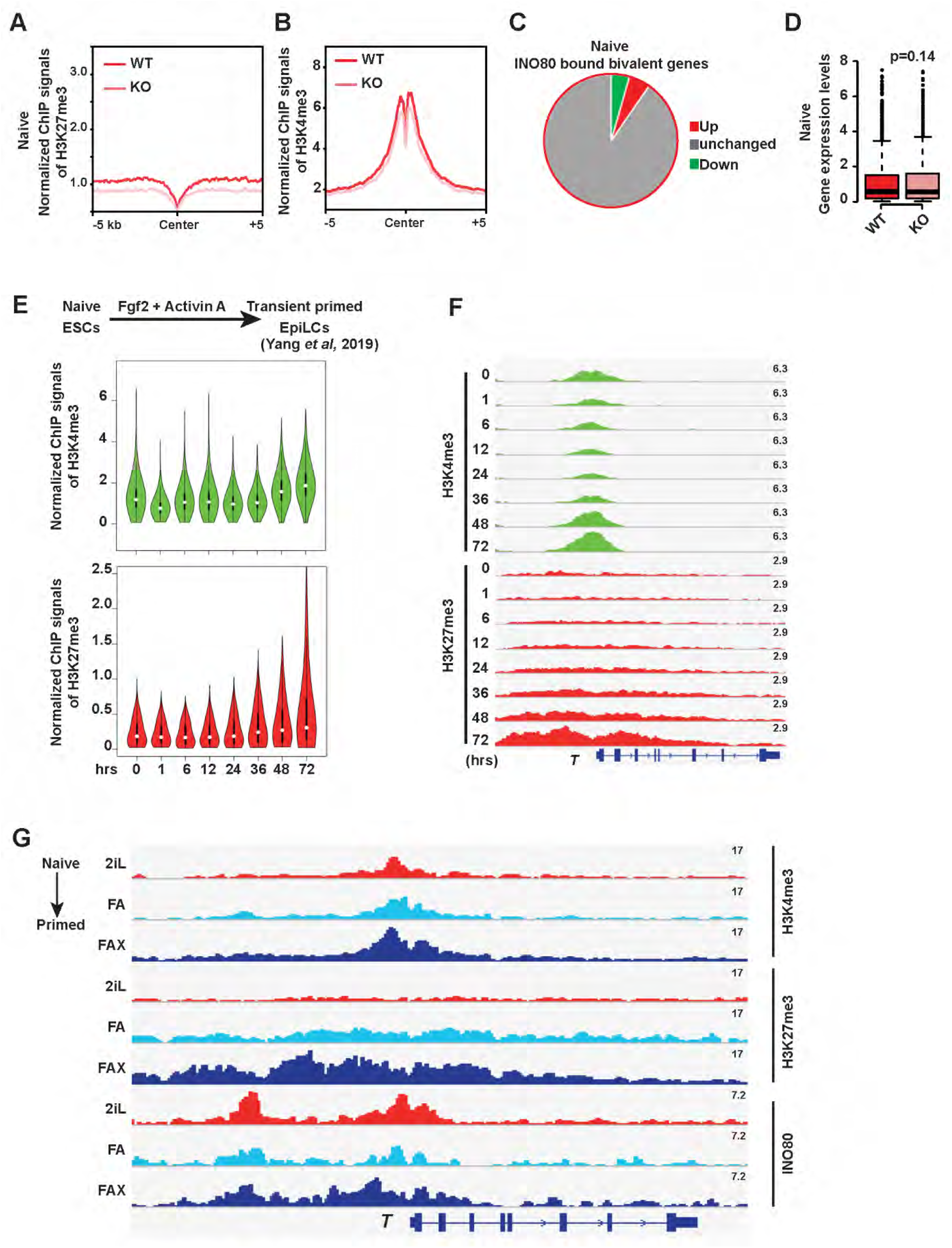
Establishment of bivalency during the naïve to primed transition (related to Figure 3–4) **(A-B)** Metagene plots to show normalized H3K27me3 (A) and H3K4me3 (B) ChIP-seq signals at INO80-bound bivalent TSSs in the naïve state. **(C-D)** Gene expression changes of INO80-bound bivalent genes after *Ino80* deletion in the naïve state. (C) Pie chart to show the number of DEGs in INO80-bound bivalent genes. (D). Box plot to show the extent of expression changes of the INO80-bound bivalent genes. p-value was calculated by Wilcoxon assigned rank test. **(E)** Violin plot to show normalized ChIP-seq signals of H3K4me3 (upper) and H3K27me3 (lower) on the primed bivalent promoters during the naïve to primed transition (based on public data). **(F)** Genome browser track to show H3K4me3 and H3K27me3 occupancy near T during the naïve to primed transition (based on public data). **(G)** Genome browser track to show INO80, H3K4me3 and H3K27me3 occupancy near T during the naïve to primed transition (from this study).

**Supplementary Figure 4.**
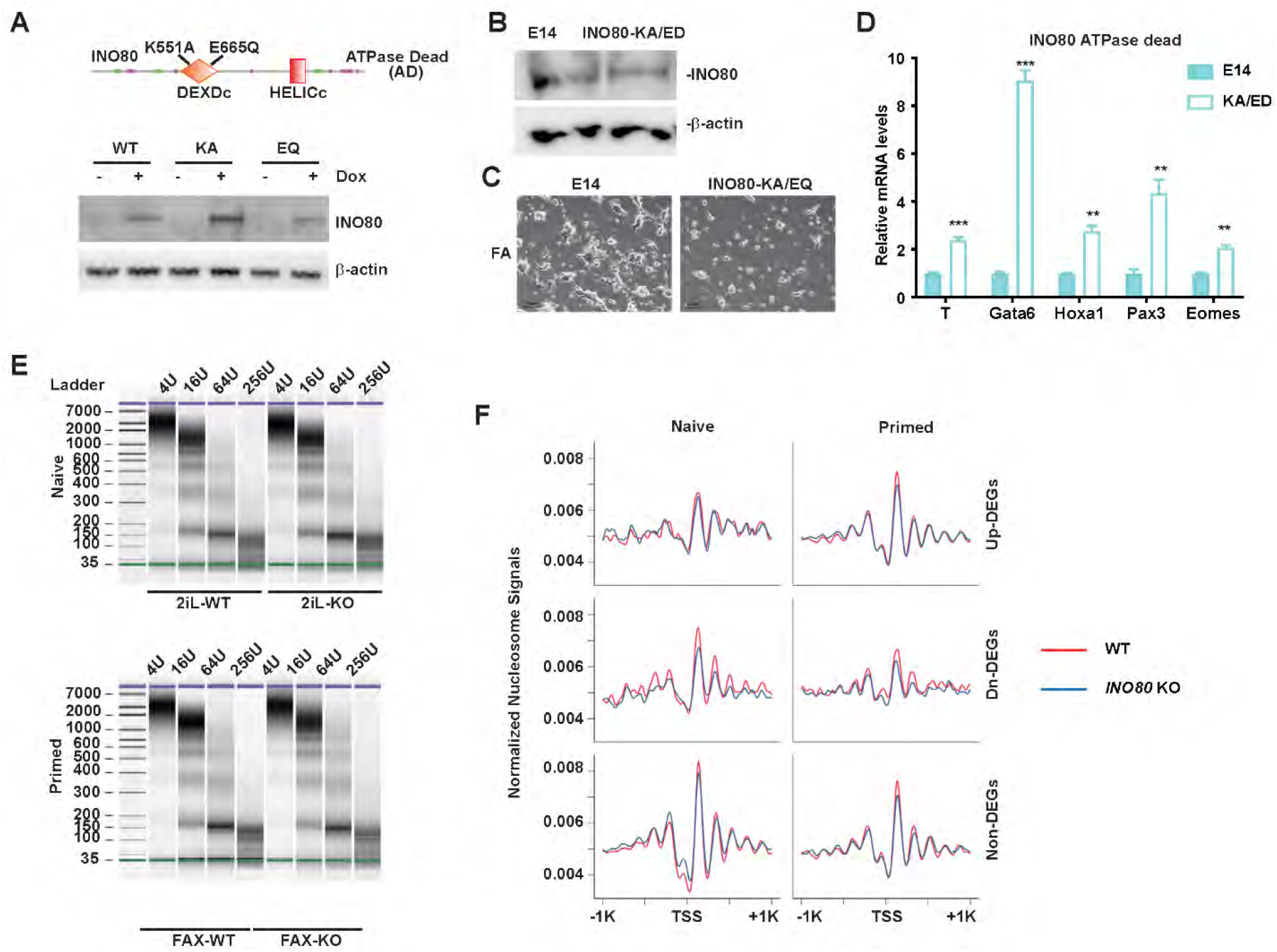
The requirement for INO80 chromatin remodeling activity (related to Figure 5) **(A)** Rescue of *Ino80* deletion phenotype by INO80-ATPase-dead mutants. *Ino80* deletion ESCs were transfected with vectors expressing Dox-inducible WT or ATPase-dead *Ino80* (KA or EQ mutant). Cells were drug selected and cultured in FAX. *Ino80* deletion was induced by 4-OHT treatment and exogenous *Ino80* expression was induced by Dox treatment. Cell morphology was imaged 4 days later. Western blot to show the expression of the Dox-inducible WT and ATPase-dead *Ino80*. **(B)** *Ino80*-KA/ED knock-in ESCs. ATPase-dead mutations were introduced into the endogenous *Ino80* in E14Tg2a cells by CRISPR-mediated genome editing. Protein expression of the WT and mutant *Ino80* was determined by western blot in WT and *Ino80* homozygous knock-in cells. **(C)** WT and *Ino80*-KA/ED knock-in ESCs were cultured in FA **(D)**. The relative expression of representative primed bivalent genes in WT and Ino80 mutation cells was determined by RT-qPCR, normalized to *Gapdh* and plotted as mean ± SEM (E). p-values were calculated by student t-test: ** <0.01, *** <0.001. **(E)** Bioanalyzer gel image to show titrated MNase digestion of WT and *Ino80* deletion cells cultured in the naïve (2iL) and primed (FAX) state. **(F)** Metagene plot of nucleosome signals at TSSs of *Ino80*-deletion induced DEGs and non-DEGs in the naïve and primed state. DEGs were separated into up-and down-regulated genes.

**Supplementary Figure 5.**
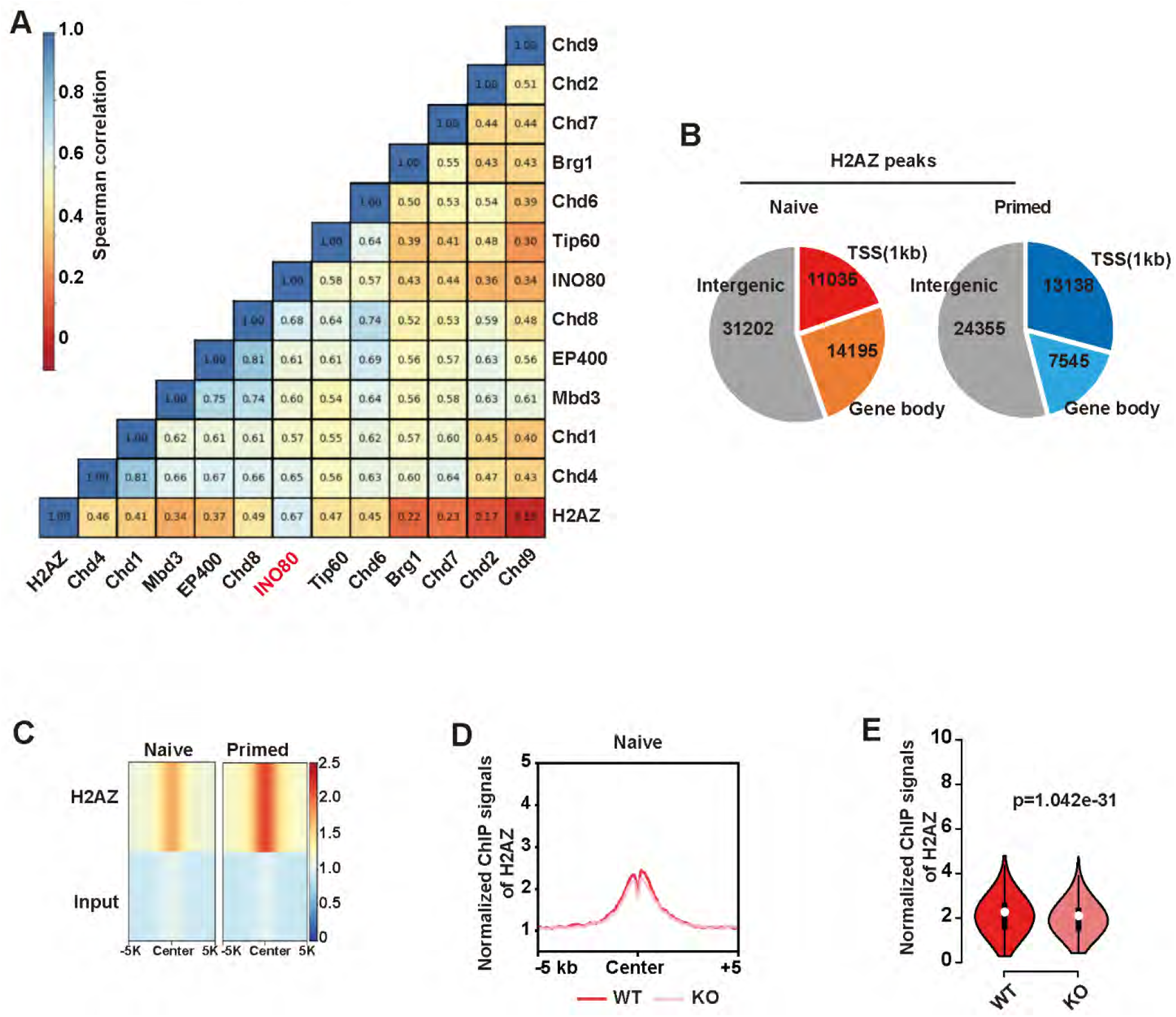
INO80 in H2A.Z occupancy (related to Figure 6) **(A)** Spearman correlation of the genomic occupancy of H2A.Z and chromatin remodelers. ChIP-seq data for INO80 and H2A.Z was from this study. ChIP-seq data for the other chromatin remodelers was from the literature. **(B)** H2A.Z ChIP-seq peak distribution in the genome in the naïve and primed state. **(C)** H2A.Z ChIP-seq signals in the naïve and primed state. **(D-E)** Metagene (D) and violin plot (E) of normalized H2A.Z ChIP-seq signals in the naïve state in WT and *Ino80* deletion cells. p-value was calculated by Wilcoxon assigned rank test.

**Supplementary Figure 6.**
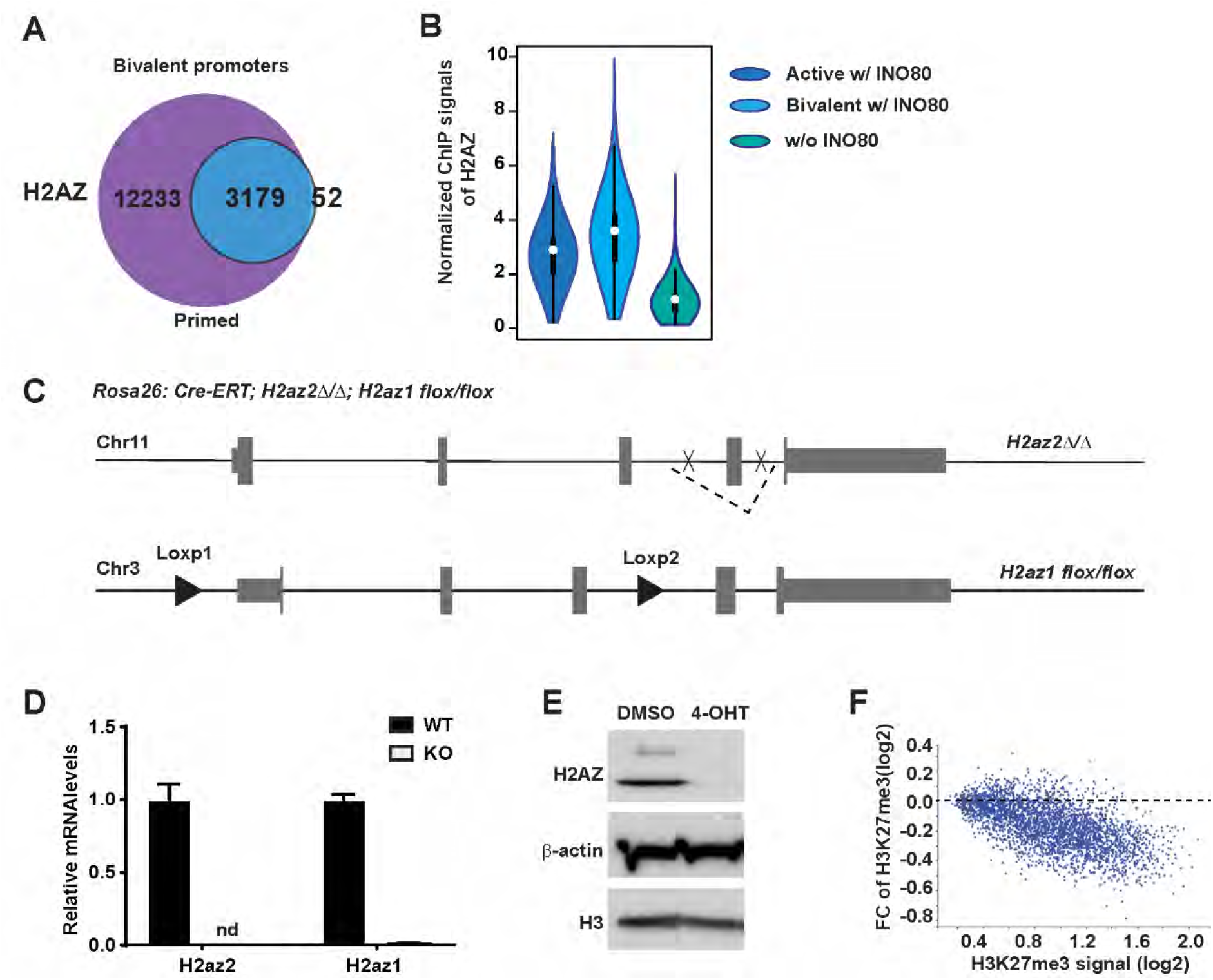
H2A.Z and bivalency (related to Figure 7) **(A)** Overlap between INO80-bound bivalent TSSs and H2A.Z peaks in the primed state. **(B)** Violin plot of normalized H2A.Z ChIP-seq signals at promoters of INO80-bound active, INO80-bound bivalent and non-INO80-bound gene promoters in the primed state. **(C)** Strategy of *H2az1/H2az2* inducible deletion in Rosa26::Cre-ERT2 ESCs. **(D)** Expression of H2A.Z in WT and *H2az1/H2az2* deletion ESCs. *H2az1*-cKO/*H2az2*-KO ESCs were treated with DMSO or 4-OHT. H2A.Z mRNA expression was determined by RT-qPCR, normalized by *Gapdh* and plotted as mean ± SEM. **(E)** H2A.Z protein expression was determined by western blot. β-actin and histone H3 were used as loading controls. **(F)** MA analysis to show changes in H3K27me3 occupancy at INO80-bound bivalent TSSs in WT and *H2az1*-cKO/*H2az2*-KO ESCs.

**Supplementary Figure 7.**
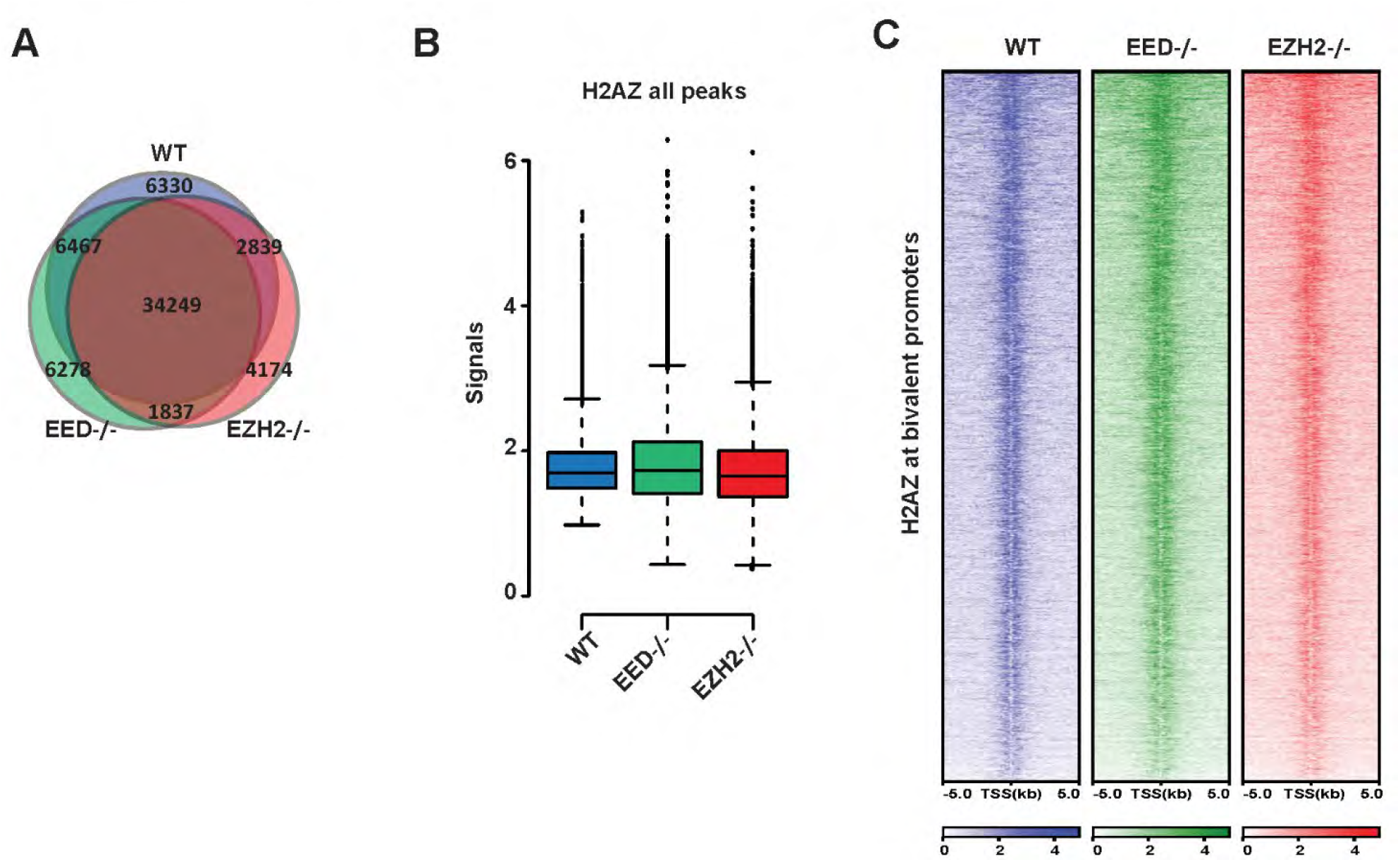
H2A.Z in PRC2 component deletion cells (related to Figure 7) **(A)** Venn diagram to show the overlap of H2AZ peaks in WT, *Eed*-KO and *Ezh2*-KO ESCs. Cells were cultured in FAX and collected for H2A.Z ChIP-seq. **(B)** Box plots to show H2A.Z ChIP-seq signals of all H2A.Z peaks in (A). **(C)** Heatmap of H2A.Z ChIP-seq signals at the primed bivalent promoters in WT, *Eed-KO* and *Ezh2*-KO ESCs.

**Supplementary Tabel S1:** Antibodies used in this study.

**Supplementary Tabel S2: Antibodies used in this study.**
|  | WB | ChIP-seq | Vendor | Cat# |
| --- | --- | --- | --- | --- |
| H3 | √ |  | Abcam | ab1791 |
| Ran | √ |  | BD | 610341 |
| b-actin | √ |  | Sigma | A5316 |
| INO80 | √ | √ | ProteinTech | 18810-1-AP |
| HA | √ | √ | Cell Signaling | #3724 |
| H2AZ | √ | √ | Active motif | 39943 |
| EZH2 |  | √ | Cell Signaling | #5246 |
| SUZ12 |  | √ | Cell Signaling | #3737 |
| H3K4me3 |  | √ | Active motif | 39159 |
| H3K27me3 |  | √ | Active motif | 39155 |
| H3K27ac |  | √ | Abcam | ab4729 |

**Supplementary Table 2:**
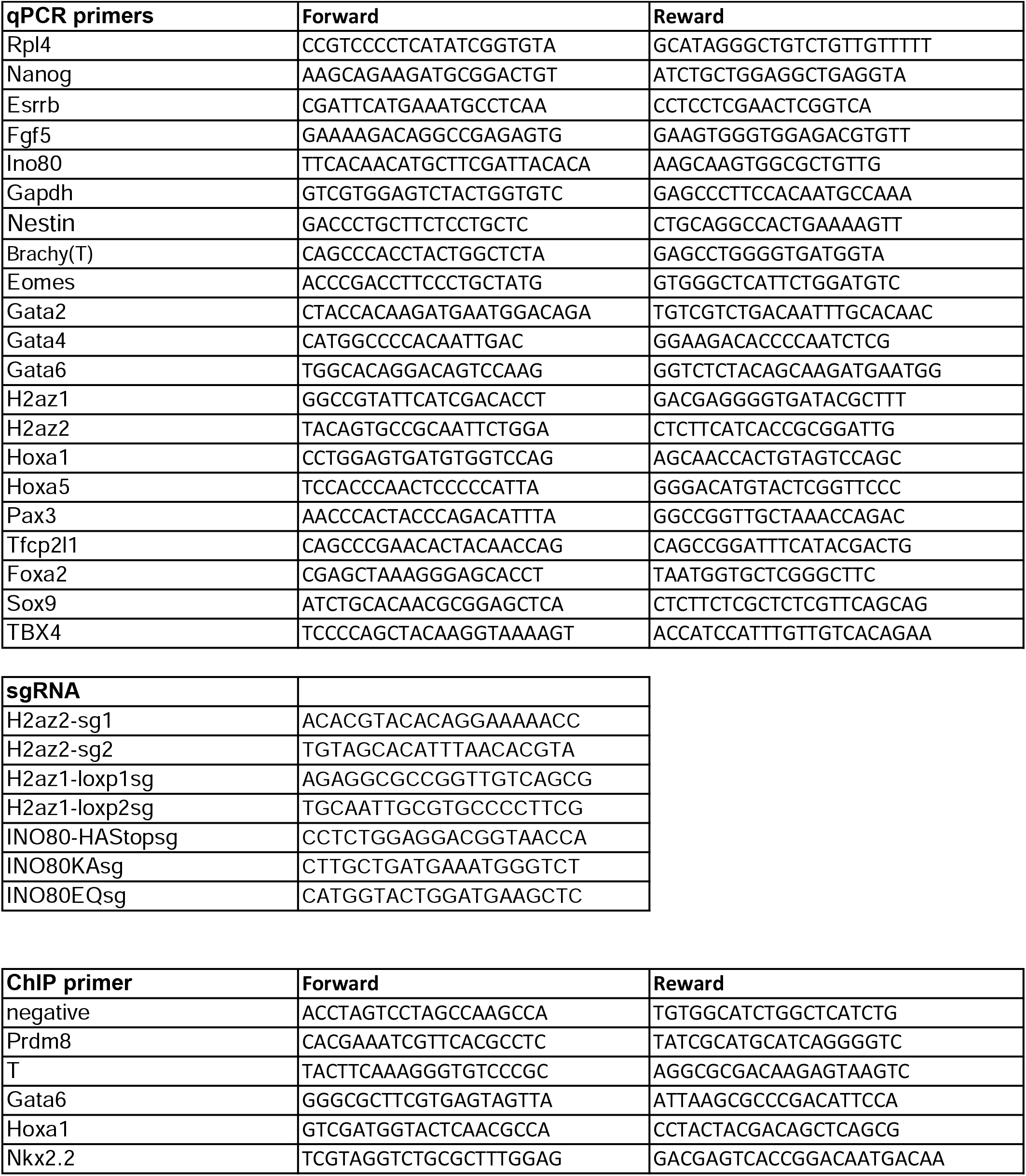
Primers for RT-qPCR, ChIP-qPCR and oligos for genome targeting.

**Supplementary Table 3.** Software and Algorithms

|  |  |  |
| --- | --- | --- |
| Bowtie (v1.1.2) | (Langmead et al., 2009) | http://bowtie-bio.sourceforge.net/bowtie2/index.shtml |
| Cutadapt (v1.9) | (Martin, 2011) | https://cutadapt.readthedocs.io/en/v1.9/installation.html |
| Samtools | (Li et al., 2009) | http://samtools.sourceforge.net/ |
| Bedtools (v2.21.0) | (Quinlan, 2014) | https://bedtools.readthedocs.io/en/latest/ |
| Deeptools | (Ramirez et al., 2016) | https://deeptools.readthedocs.io/en/develop/ |
| STAR (v2.5.2b) | (Dobin et al., 2013) | https://github.com/alexdobin/STAR |
| DESeq2 1.16.1 | (Love et al., 2014) | https://bioconductor.org/packages/release/bioc/html/DESeq2.html |
| normalize.loess R affy package | (Cleveland, 1992; Gautier et al., 2004) | https://www.rdocumentation.org/packages/affy/versions/1.50.0/topics/normalize.loess |
| bedGraphToBigWig | (Kent et al., 2010) | https://www.encodeproject.org/software/bedgraphtobigwig/ |
| R 3.3.0 | R Development Core Team, 2016 | https://www.r-project.org/ |
| BoxplotR | (Spitzer et al., 2014) | https://github.com/VizWizard/BoxPlotR.shiny/blob/master/README.md |

